# Functional associations of evolutionarily recent human genes exhibit sensitivity to the 3D genome landscape and disease

**DOI:** 10.1101/2024.03.17.585403

**Authors:** Katherine Fleck, Victor Luria, Nitanta Garag, Amir Karger, Trevor Hunter, Daniel Marten, William Phu, Kee-Myoung Nam, Nenad Sestan, Anne H. O’Donnell-Luria, Jelena Erceg

## Abstract

Genome organization is intricately tied to regulating genes and associated cell fate decisions. Here, we examine the positioning and functional significance of human genes, grouped by their lineage restriction level, within the 3D organization of the genome. We reveal that genes of different lineage restriction levels have distinct positioning relationships with both domains and loop anchors, and remarkably consistent relationships with boundaries across cell types. While the functional associations of each group of genes are primarily cell type-specific, associations of conserved genes maintain greater stability across 3D genomic features and disease than recently evolved genes. Furthermore, the expression of these genes across various tissues follows an evolutionary progression, such that RNA levels increase from young lineage restricted genes to ancient genes present in most species. Thus, the distinct relationships of gene evolutionary age, function, and positioning within 3D genomic features contribute to tissue-specific gene regulation in development and disease.

## Introduction

The genome is intricately folded and regulated in the nucleus, which may contribute to cell type diversity. Recent technological advances in mapping of chromosomal interactions and single-cell imaging have provided insights into the organization of chromatin at various levels^1–4^. Domains (also termed topologically associating domains or TADs) with increased chromatin interactions may be insulated from neighboring regions by boundaries with decreased chromatin interactions^5–8^. Formation of these domains is often mediated by extrusion of chromatin loops, which dynamically link distal anchor points ^6,9–12^. However, the interplay between genome structure and function is still unclear^13–16^. For instance, while some chromosomal disruptions and perturbations of regulators involved in genome folding may not lead to substantial effects on gene regulation, other structural disruptions at specific loci may affect contacts between regulatory regions, leading to cancer and congenital limb malformations^13,14,17–22^. Moreover, communication between regulatory regions associated with gene activation is facilitated by direct chromatin looping or other hub-like structures^6,13,14,23–34^. Having proper regulatory interactions, including those between enhancers and promoters, is key for cell type-specific gene activity.

Understanding how the genome is organized and the extent to which its organization is conserved may illuminate the functional significance of genomic regions. For instance, perfectly conserved sequences between distantly related species that diverged 90 to 300 million years ago, named ultraconserved elements (UCEs)^35–38^, have non-random relationships to 3D genome organization^39^. Specifically, UCEs are enriched in domains and associated with renal and neuronal processes when present in domains common to many cell types^39^. Clusters of conserved noncoding elements (CNEs) around developmental genes, with 70-90% conservation level, are also highly concordant with domains^40^. Moreover, human accelerated regions (HARs), which are sequences conserved in vertebrates but fast-evolving in the human lineage^41–49^, are associated with domains that have human-specific structural variants, suggesting potential rewiring of contacts between regulatory regions^50^. Moreover, structural variants and mutations affecting conserved and fast-evolving regions are often related to diseases and neurodevelopmental disorders^37,46,51–55^.

Protein-coding genes also experience varying levels of conservation, from extremely high sequence conservation across evolutionary taxa to recent evolution in specific lineages. New protein-coding genes originating by non-copying mechanisms (for example, *de novo* from intergenic regions; from long non-coding RNAs that become coding; by overprinting, that is, translating an existing mRNA in a different frame; or by translating an antisense RNA^56,57^) have been discovered by studying naturally occurring gene variants, through mutagenesis screens, and by proteogenomics in species ranging from plants to humans^58–64^. New genes encoding sequences not similar to existing proteins are expressed in different tissues^65,66^, have function^59,67–71^ and are formally defined as lineage restricted genes (LRGs). In contrast to new genes clearly arising from copying mechanisms (gene or genome duplication; retrotransposition; exon shuffling; or pseudogene resurrection), LRGs can have new protein domains^70–72^. To understand how gene expression and function emerge and change with time in the context of a continuous process of gene birth and death^73^, genes of different evolutionary ages can be compared once the minimal evolutionary age of each gene is evaluated by sequence similarity or synteny^58,74,75^.

Here, we asked how the positionings of genes of different evolutionary ages in the 3D genome landscape relate to their expression and functional significance. Interestingly, while genes of different evolutionary ages have distinct relationships with domains and loop anchors, their relationships with boundaries are consistent across cell types. Furthermore, evolutionarily older genes maintain stable functional associations regardless of genomic positioning and are expressed at higher levels than more recently evolved genes, which exhibit functional associations that are sensitive to the positioning of these genes within the 3D genome organization and disease state. Together, the distinct 3D genomic positionings of genes as a function of their evolutionary age may be related to the regulation of tissue-specific programs in development and disease.

## Results

### Estimation of gene evolutionary age more conservative by sequence similarity than by synteny

To understand the relationship between gene age and features of 3D genome organization, we built a dataset with all human protein-coding genes and control intergenic sequences. First, we evaluated the evolutionary age of all human protein-coding genes (Figures 1A, S1A, and S1B). Of these 21,436 genes (Table S1A, to avoid double counting, for each protein-coding gene, we determined the age of each isoform, and selected the longest of the oldest isoforms), most (19,334) are annotated in Ensembl 89 (Figure S1A). Additionally, 2,102 unannotated genes with translation evidence from proteogenomics studies^76–79^ were included in the analysis (Figure S1B; Table S1A). We estimated gene evolutionary age by evaluating lineage restriction level using protein sequence similarity. This method estimates the minimal evolutionary age of a gene by determining its level of taxonomic lineage restriction, which is indicated by the most distant species in which a protein sequence sufficiently similar to the query sequence is detected, and inferring that the gene must be at least as ancient as the common ancestor of the query species and the most distant species^74,80,81^. The method estimates gene age as the age of the most ancient part of the protein. The youngest genes identified by this method are the ones detectable in the fewest species, or just one species, human in this analysis. We refer to these genes as *lineage restricted genes* (LRGs). The oldest genes are the ones detectable in all species. Thus, a human gene for which a sufficiently similar sequence exists in bacteria is ancient (14,845 genes); a human gene also present in sponges but not in more distant species is specific to metazoans (2,253 genes); a human gene also present in lancelets but not in more distant species is specific to chordates (1,584 genes); a human gene also present in platypus but not in more distant species is specific to mammals (685 genes); and a human gene also present in lemurs but not in more distant species is specific to primates (2,069 genes) (Figure 1B). We refer to these five categories (ancient, metazoan, chordate, mammal, primate) as *eras*, and to genes within them as *era genes*. As controls for human protein-coding genes of varying evolutionary age, we used intergenic open reading frames (Igen ORFs, 3,924 sequences, Table S1B) that are potential protein-coding sequences with no known evidence of transcription or translation. Additionally, we used human intergenic sequences that cannot encode proteins (Igen Non-ORFs, 3,892 sequences, Table S1C) and are matched in length to Igen ORFs. We find that there is an evolutionary progression from short young genes to long ancient genes (Figures 1C and 1D), as noted previously in multiple eukaryotic species^58,74,82^. Interestingly, the youngest genes are very similar in size to control Igen ORFs (Figures S1C and S1D), suggesting that young genes are picked at random from the myriad of ORFs present in the vast intergenic parts of the human genome.

**Figure 1.**
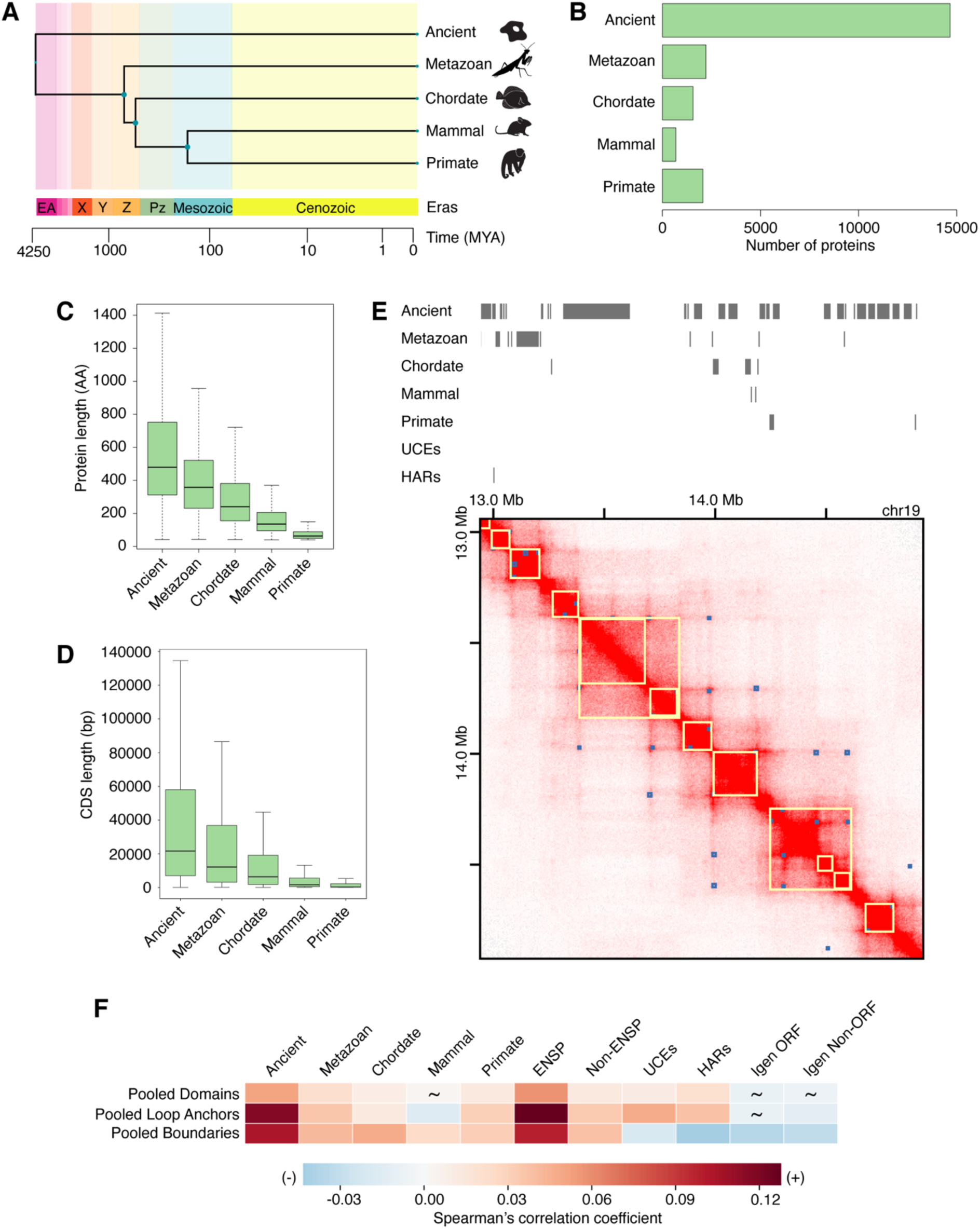
Genomic regions of varying sequence conservation have distinct relationships with features of 3D genome organization. (A) Human genes assigned an evolutionary age by phylogenomics and phylostratigraphy. Representative species used with TimeTree^149^ to build a timetree: *Escherichia coli* (ancient), *Amphimedon queenslandica* (metazoan), *Branchiostoma floridae* (chordate), *Ornithorhynchus anatinus* (mammal), and *Homo sapiens* (primate). (B) Number of human protein-coding genes in every evolutionary era (ancient, metazoan, chordate, mammal, primate). Most genes are ancient. (C) Protein length is indicated in number of amino acids (AA) and is largest for ancient genes. (D) The length of coding sequences (CDS) is indicated as the number of base pairs (bp) and is largest for ancient genes. (E) Visualization of example genomic regions of varying sequence conservation with the GM12878 Hi-C dataset^6^ using the Juicebox tool^92^. Domains (yellow squares) and peaks indicating loop presence (blue squares), as annotated in Juicebox. (F) Pooled domains, loop anchors, and boundaries are significantly positively correlated with era genes (1.08×10^-172^ ≤ p ≤ 0.022), except for mammal genes in pooled domains and loop anchors (6.03×10^-5^ ≤ p ≤ 0.654). UCEs and HARs are significantly positively correlated with pooled domains and loop anchors, but negatively correlated with pooled boundaries (1.34×10^-29^ ≤ p ≤ 0.017). Spearman correlation analysis was performed by partitioning the genome into 50-kb bins. Spearman correlation coefficients are indicated by a heatmap. Non-significant p values are depicted by tildes. Control datasets include annotated genes, unannotated genes, Igen ORFs, and Igen Non-ORFs.

Within LRGs, a subset can be ascertained to have arisen *de novo* if a similar non-coding sequence is found in a syntenic region of the genome of a taxonomically close species. Loss of synteny driven by recombination limits such ascertainments to genes that appeared in the last 100-200 million years^56,74^ and thus excludes the majority of genes, which are situated in genomic regions that are not syntenic between distant species. Therefore, to study the majority of protein-coding genes, we focus on lineage restriction evaluated by sequence similarity. This is a valuable tool to estimate the minimal age of the gene because the probability of independent emergence of two similar proteins decreases exponentially with protein length. Since synteny and sequence similarity are independent tools, we compared the results of gene age estimation by synteny^83^ versus by sequence similarity^80–82^ using 18,098 human genes that are common to GenTree^83^ and to our list of human genes. First, we found that many genes appear considerably younger by synteny than by sequence similarity, as only 60.0% of genes (10,867/18,098) are early genes by synteny versus 94.8% (17,154/18,098) by sequence similarity. Thus, the younger evolutionary branches contain more genes by synteny than by sequence similarity (Figure S1E; Table S1D). Second, for every gene, we compared the gene ages from the two methods and found that, while for 61.2% of the genes there is no difference (11,075/18,098), most of the remaining genes appear younger by synteny (37.1%, 6,707/18,098), and only very few (1.8%, 316/18,098) appear younger by sequence similarity (Figure S1F; Table S1D). These observations indicate that sequence similarity provides gene age and taxonomic restriction estimates that are more conservative than those obtained from synteny, consistent with the criteria used by each method. Synteny evaluates the taxonomic restriction of a gene by examining its genomic context defined by the identity of the closest genes, while sequence similarity evaluates the taxonomic restriction of the oldest protein-coding sequence interval of the gene. Thus, a gene recently duplicated from an ancient gene, such as NOTCH2NL^84,85^ and ARHGAP11B^86^, or copied and retrotransposed to a genomic location different from that of the parent gene^87,88^, will appear recent by synteny but older by sequence similarity. While chromosomal context matters to the regulation of the expression of individual genes and is the focus of synteny methods^75,83,89,90^, the alternative point of view, which our work prioritizes, is that properties of the protein-coding regions matter to gene function, and are thus the focus of protein sequence similarity methods. Furthermore, as we showed here, estimates of evolutionary age are more conservative from sequence similarity than from synteny.

Second, we included two datasets with different levels of sequence conservation, but comparable numbers of elements to the era genes. The first dataset is comprised of UCEs, genomic regions at least 200 base pairs (bp) in length that share 100% sequence identity between reference genomes of distantly related species. Comparisons between human, mouse, and rat; or human, dog, and mouse; or human and chicken genomes identified 896 UCEs in total^39,53,91^. The second dataset is comprised of >3,100 HARs which are sequences that are conserved in vertebrates but are rapidly evolving in the human genome^41–49^. Both of these regions may be often non-protein coding, and many have been associated with regulatory functions. As such, they may distribute differently from protein-coding genes in the 3D genomic context.

### Regions of varying sequence conservation have distinct relationships with 3D genomic features across cell types

We began by visualizing the placement of era genes, UCEs, and HARs in the 3D genome landscape, as illustrated with an example genomic region of the GM12878 cell line^6,92^ (Figure 1E). Then, to globally assess how genomic regions of varying sequence conservation relate to features of 3D genome organization such as domains, loop anchors, and boundaries, we compiled Hi-C datasets from various published studies encompassing a wide range of cell types (Table S1E)^5,6,19,93–103^. Due to the variation in methodology between these studies (e.g. Hi-C protocol, sequencing depth, quantity of input material), we considered each dataset individually as well as grouped by 3D genomic feature to generate pooled datasets. Then, we performed correlation analyses as previously described^39,53^. Briefly, the genome was partitioned into equal bins of defined size, within which the proportion of sequence covered by each feature of interest was determined. The global pairwise correlations between feature densities were performed to generate Spearman correlation coefficients and p values. These analyses were carried out using a breadth of bin sizes (20 kb, 50 kb, and 100 kb) to account for the differences among individual 3D genomic features in terms of their number of regions, median region size, and proportion of genome covered (Table S1E).

We assembled 18 domain datasets from six studies^5,6,93,95,96,103^ covering 14 human cell types, including human embryonic and induced pluripotent stem cells as well as fetal cells, differentiated cells, and cancer cell lines representing different germ layers (Table S1E). Individually, these domain datasets cover varying percentages of the genome, ranging from 40.07% (HMEC) to 87.72% (SK-N-SH). Pooling the datasets together and collapsing the overlapping intervals resulted in a single dataset (hereafter named “pooled domains”) spanning 98.06% of the genome (Table S1E). Pooled domains show predominantly significant, positive correlation with genes of most evolutionary eras, UCEs, and HARs (6.27×10^-35^ ≤ p ≤ 0.022), except mammal genes (p = 0.654) (Figure 1F; Table S2). When examined by individual cell types, ancient genes and UCEs continue to display mainly significant, positive correlations with domains (0 ≤ p ≤ 0.746; Figure S2A; Table S2). This observation aligns with previous findings that UCEs and clusters of CNEs, which may have regulatory function, coincide with domains^39,40^. Notably, annotated genes also closely match the trend in correlations between ancient genes and domains (0 ≤ p ≤ 0.078; Figure S2A; Table S2). Interestingly, less conserved era genes (metazoan, chordate, mammal, and primate) and fast-evolving HARs have more variable strengths of correlations with domains across cell types (2.34×10^-92^ ≤ p ≤ 0.988; Figure S2A and Table S2). Taken together, these observations suggest that conserved elements have more consistent relationships with domains, while less conserved elements show more variable relationships with domains across cell types.

We continued our analyses using 36 datasets of loop anchors from various studies^6,19,93,94,97–102^. These datasets span 36 distinct cell types ranging from H1 human embryonic stem (ES) cells, ES-cell-derived lineages, fetal cells, to differentiated and cancer cell lines (Table S1E). Loop anchors cover a smaller portion of the genome than domains; these individual datasets range from 1.97% (RPE1) to 9.83% (aorta). Again, we pooled the loop anchor datasets together and collapsed the overlapping intervals to generate a dataset named “pooled loop anchors,” which covers 56.55% of the genome (Table S1E). Similarly to pooled domains, pooled loop anchors display mostly significant, positive correlations with genes of most eras, UCEs, and HARs (1.08×10^-172^ ≤ p ≤ 0.003), except for mammal genes, whose correlation is significantly negative (p = 6.03×10^-5^) (Figure 1F; Table S2). Notably, HARs have previously shown enrichment in loops specific to human compared to rhesus macaque and mouse^104^. While the observed significant, positive correlation in UCEs differs from a former study^39^, which displayed a non-significant relationship between UCEs and loop anchors, this result can be explained by the increase in the number of datasets used in this analysis (36 datasets here compared to previously used eight datasets). Our observation that UCEs are positively correlated with pooled loop anchors is consistent with the regulatory activity of some UCEs^105–113^ and with loops facilitating enhancer-promoter contacts^6^. When separated by cell type, UCEs together with HARs and less conserved era genes (metazoan, chordate, mammal, primate) have variable relationships with loop anchors (8.65×10^-118^ ≤ p ≤ 0.965; Figure S2B; Table S2). On the other hand, ancient and annotated genes show a robust, significant positive correlation with loop anchors across most cell types (0 ≤ p ≤ 0.944; Figure S2B; Table S2). In addition, we noted that relationships of genes of different eras and HARs with loop anchors remain persistent across some cell types, mostly corresponding to Hi-C datasets produced with the same methodology (Figure S2B; Tables S1E and S2). Overall, ancient genes have the most consistent relationships, while younger LRGs, HARs, and UCEs have more variable relationships with loop anchors across cell types.

Finally, we turned to boundary datasets from multiple studies^5,94,98,102^. These boundary datasets cover 20 cell types and primary tissues (Table S1E). Individual datasets cover portions of the genome more comparable to loop anchors than domains ranging from 2.61% (aorta) to 3.95% (hESC). A dataset of pooled boundaries covers 22.5% of the genome (Table S1E). This pooled dataset is significantly positively correlated with genes of different eras (2.62×10^-136^ ≤ p ≤ 2.90×10^-9^), consistent with transcription start sites being enriched at boundaries^5^. Conversely, pooled boundaries are significantly negatively correlated with control Igen ORFs (p = 9.06×10^-18^) and Igen Non-ORFs (p = 2.11×10^-14^), UCEs (p = 1.81×10^-6^), and HARs (p = 1.76×10^-25^) (Figure 1F; Table S2). These observations in UCEs and HARs are different from those found with pooled domains and loop anchors (Figure 1F) and are consistent with the depletion of UCEs at boundaries^39^. Furthermore, in distinct contrast to domains and loop anchors, genes of different eras, UCEs, and HARs all show remarkable consistency in their correlations with boundaries across cell types (Figure S2C; Table S2). Strikingly, genes display predominantly significant, positive correlations with boundaries (2.19×10^-110^ ≤ p ≤ 0.306), as opposed to HARs with entirely significant, negative correlations (8.91×10^-12^ ≤ p ≤ 0.009) and UCEs with significant or non-significant negative correlations (6.24×10^-4^ ≤ p ≤ 0.136) (Figure S2C; Table S2).

Additional correlation analyses with different bin sizes (20 and 100 kb) provide similar results to 50-kb results for all three 3D genomic features (Table S2). Moreover, the control sets of Igen ORFs and Igen Non-ORFs, which have no evidence of regulated transcription, support these findings with era genes by displaying similar correlations regardless of the inspected 3D genomic feature or cell type (Figures 1 and S2; Table S2). Taken together, the relationships between genomic regions of varying sequence conservation and boundaries are consistent across cell types yet distinct between era genes and UCEs and HARs, while the relationships with domains and loop anchors are variable across cell types. This variability, which is more prominent with evolutionarily younger genes and fast-evolving HARs, may suggest these regions’ potential role in tissue-specific programs.

### Gene expression changes with evolutionary age

Due to the interplay between the 3D organization of the genome and gene expression, we analyzed the transcription of genes and intergenic control sequences using the Genotype to Tissue Expression (GTEx) database^114^, which contains RNA sequencing data from 54 human adult tissues (Tables S3A and S3C; S3B and S3D for significance testing). We grouped the data into six categories: the three germ layers (ectoderm, mesoderm and endoderm), the germ line (ovary, testis), and the brain, since some evolutionarily novel genes function in neurons^59,65,70,115^. First, we examined the RNA expression of genes (21,436 genes: 14,845 ancient, 2,253 metazoan, 1,584 chordate, 685 mammal, 2,069 primate; 19,334 annotated genes, 2,102 unannotated genes) as a function of their evolutionary age (Figures 2 and S3A; Tables S3A and S3E). We found that gene expression changes with evolutionary era, such that, in all tissue categories, ancient genes are expressed at the highest levels while primate genes are expressed at significantly lower levels (Tables S3A and S3B; for example, RNA Mean Counts in Ectoderm: Ancient: 2,709 ± 56, 1,182, from 14,845 genes, versus Primate: 118 ± 17, 13, from 2,069 genes, average ± standard error of the mean, median, number of genes; p < 2.225×10^-308^). Moreover, the expression of all genes is 100-1000 times higher than that of control Igen ORFs and Igen Non-ORFs, which reflect the transcriptional background of the genome (Tables S3A and S3B; for example, Ectoderm: Primate: 118 ± 17, 13, from 2,069 genes, versus Igen Non-ORF: 0.23 ± 0.04, 0.006, from 3,889 sequences, average ± standard error of the mean, median; p < 2.225×10^-308^).

**Figure 2.**
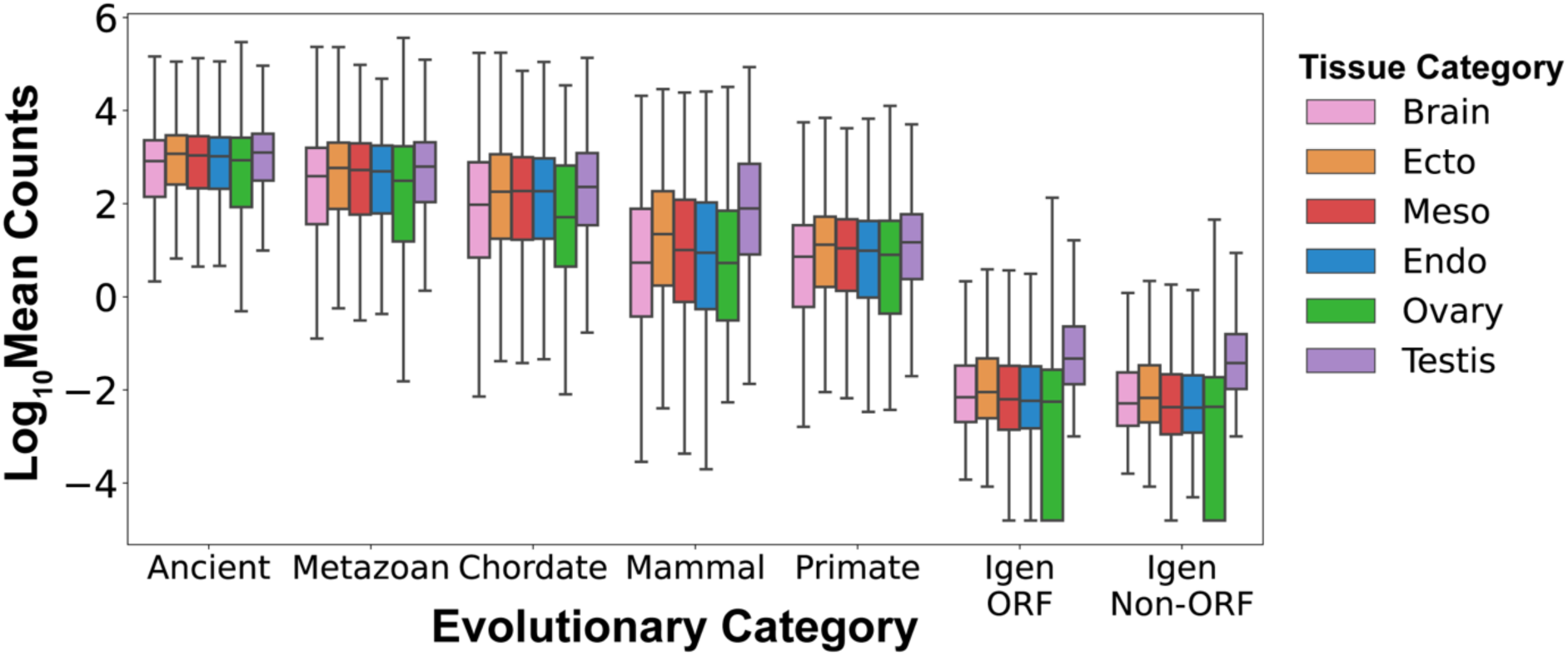
Transcriptional levels vary with gene age and tissue of origin. The expression of human protein-coding genes is measured as the log10 of the mean counts of RNA transcripts across 54 human tissues from the GTEx database^114^. RNA expression levels vary as a function of evolutionary category, increasing from youngest to oldest genes, and higher in all genes than in control non-genic sequences (Igen ORF, Igen Non-ORF). Testis expression is higher than expression in other tissues in control non-genic sequences.

We tested if these observations also hold if gene age is estimated by chromosomal synteny instead of protein sequence similarity. We computed the RNA expression of 18,098 human genes classified by synteny^83^ and now also by sequence similarity into four evolutionary eras (Early, Vertebrate, Mammal, Primate; Tables S3F and S3G). The era division employed here is constrained by the classification of the synteny-based GenTree database^83^, whose oldest taxonomic branch spans all taxonomic nodes from phylostratum (PS) 1, Cellular organisms, to PS 13 Euteleostomi. We found that, with both methods, early genes are expressed at higher levels than primate genes (Figures S3B and S3C; Table S3D; for example, Ectoderm by synteny: Early 3,093 ± 64, 1,597, from 10,867 genes, versus Primate 499 ± 223, 9, from 684 genes, average ± standard error of the mean, median, number of genes; p < 2.225×10^-308^; Ectoderm by sequence similarity: Early 2,585 ± 52, 1,079, from 17,154 genes, versus Primate 676 ± 171, 32, from 142 genes, average ± standard error of the mean, median, number of genes; p < 2.225×10^-308^). While a substantial fraction of human genes (37%) appear younger by synteny, the RNA expression levels of primate genes are in the same range for both synteny and sequence similarity (Figures S3B and S3C; Table S3D; Ectoderm by synteny 499 ± 223, 9 from 684 genes, versus 676 ± 171, 32 from 142 genes, by sequence similarity; p = 0.003) while early genes have higher expression levels by synteny (for example, Ectoderm by synteny 3,093 ± 64, 1,597, from 10,867 genes, versus 2,585 ± 53, 1,079, from 17,154 genes by sequence similarity; p = 6.55×10^-108^). Taken together, these results show that the evolutionary progression of RNA expression levels from low in young genes to high in ancient genes is robustly detected with two different gene age estimation methods.

Second, we examined the RNA expression of genes within 6 tissue categories (Figures S3D and S3E) and found that within each category there is a similar evolutionary progression of gene expression, such that ancient genes are expressed at higher levels than evolutionarily young genes. Interestingly, within the testis, the expression of genes is higher than in other tissues, as observed in other species such as fruit fly^66,116^. We observed that, for most genes of all eras, RNA expression in the brain is similar to or lower than in other tissues (Figures S3D). Within tissue categories, gene expression can vary substantially between individual tissues (Figure S4). Within the brain, ancient genes exhibit the highest RNA levels in the cerebellar hemispheres while their lowest RNA levels are in the amygdala and *substantia nigra* (Figure S4). Additionally, we observed that, while the expression of genes generally increases with gene length (short genes are expressed at lower levels than long genes), within each gene length decile the evolutionary progression from ancient genes expressed at high levels to young genes expressed at low levels is maintained (Figure S3F). Taken together, these data show that gene expression increases with evolutionary age and that all protein-coding genes are expressed at levels much higher than the transcriptional background of the genome.

### Era genes are primarily associated with prominent cell type-specific processes

Given the distinct relationships between regions of varying sequence conservation and 3D genome structure across cell types, we next investigated the functional associations of those regions. Previous work associated Ensembl-annotated genes grouped by their evolutionary age with specific gene ontology (GO) terms^117^. Due to the complexity of our datasets, which comprise recently curated annotated genes and as yet unannotated genes, we used the Genomic Regions Enrichment of Annotations Tool (GREAT)^118^ to assess GO term enrichments in genomic coordinates of interest. This approach allowed for inspection of different genomic coordinates of interest, including those of HARs, UCEs, and unannotated genes, particularly primate genes, without a need to rely on gene identifiers as inputs. We observed that metazoan genes are enriched for GO terms related to signaling pathways against all era genes as a background (Figure S5A). Meanwhile, chordate genes are associated with immune response-related processes (Figure S5B). As for mammal and primate genes, both are linked to keratinization and keratinocyte differentiation processes (Figures S5C and S5D). However, mammal genes are also associated with defense response to other organisms (Figure S5C). Ancient genes do not show enrichment in GO terms, most likely since they form a majority (69.25%) of all era genes. Together, these observations are consistent with a previous report^117^ and suggest that various categories of era genes are predominantly associated with distinct cell type-specific GO terms.

### Evolutionarily older regions show stable functional associations within 3D genomic features

Having the corresponding functional associations of era genes, we examined whether these associations are related to positioning of era genes within 3D genomic features. Expectedly, era genes in pooled domains are associated with the same GO terms (Figures 3A-3D) as with the entire genome (Figure S5). Pooling the domains across cell types may include regions that are domains in certain cell types but are boundaries and loop anchors in other cell types, and therefore incorporates more than 99% of the genes. In pooled loop anchors, metazoan and chordate genes still maintain very similar enrichment of GO terms (Figures S6A and S6B) as noted with genes in pooled domains (Figures 3A and 3B). On the other hand, evolutionarily younger mammal and primate genes lose association with defense response to other organisms and peptide cross-linking, respectively (Figures S6C and S6D). As for pooled boundaries, we observed a notable shift in the gene eras that exhibit GO term enrichment. For instance, ancient genes in pooled boundaries appear mainly associated with protein localization to telomeres as well as various ncRNA processes (Figure S6G), consistent with the enrichment of housekeeping genes at boundaries^5^. Meanwhile, metazoan and chordate genes in pooled boundaries show stable association with signaling pathways and immune response-related processes, respectively (Figures S6H and S6I). Interestingly, evolutionarily recent mammal and primate genes in pooled boundaries do not display any GO term enrichments. Overall, the functional associations of older metazoan and chordate genes seem to be stabilized and not affected by their positioning within 3D genomic features. In contrast, functional associations of younger mammal and primate genes may change depending on the 3D genomic features these genes overlap, suggesting sensitivity to their positioning within genome architecture.

**Figure 3.**
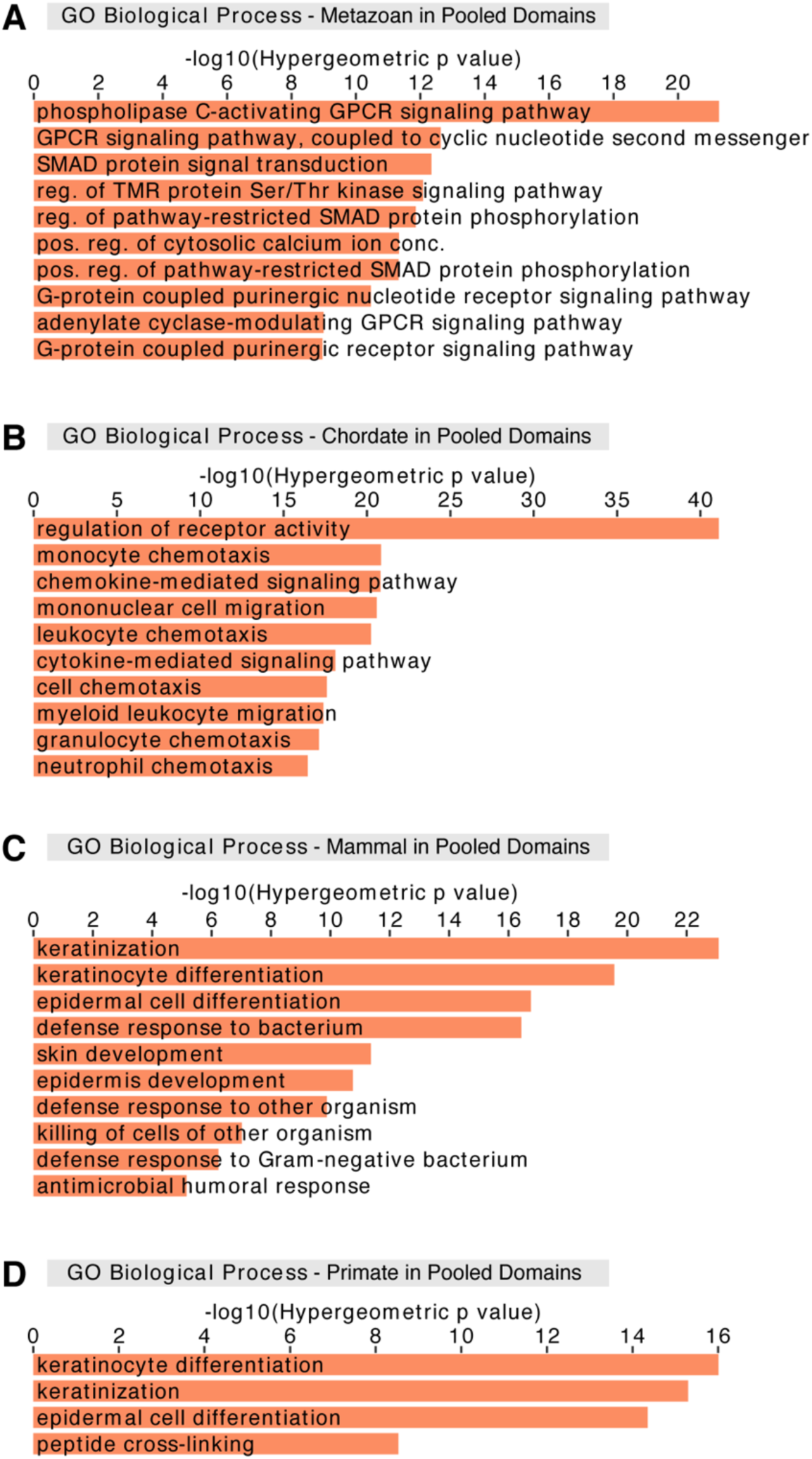
Genes of different evolutionary eras are associated with prominent cell type-specific processes in pooled domains. GREAT analysis for different era genes that overlap pooled domains with GO terms associated with (A) signaling pathways for metazoan genes, (B) immune response-related processes for chordate genes, (C) skin development and defense response to other organisms for mammal genes, and (D) keratinocyte differentiation for primate genes. (A-D) Analysis performed against all era genes as a background. Abbreviated GO terms: GPCR, G-protein coupled receptor; reg., regulation; TMR, transmembrane receptor; Ser, serine; Thr, threonine; pos., positive; conc., concentration.

We next compared our functional associations of era genes in various 3D genomic features with those of UCEs and HARs. Previous studies suggest that specific subsets of UCEs are associated with RNA processing and developmental processes^35,39,119–122^. In line with this, we observed that all UCEs in pooled domains, boundaries, or loop anchors maintain consistent stable enrichment for GO terms related to transcription and RNA metabolic processes (Figures S6E, S6J, and S16K)^35,121,123^. HARs in pooled domains, which comprise all but one HAR in our dataset, are associated with neuronal processes, cartilage formation, and thymus development (Figure S6L), consistent with functional implications of HARs due to their increased divergence in humans^42–44,46–48,50,52^. As for HARs in pooled loop anchors, associations are minorly different from those for HARs in pooled domains (Figure S6F). In pooled boundaries, HARs do not show enrichment for GO terms, which corresponds to the strong negative correlation between HARs and pooled boundaries (p = 1.76×10^-25^; Figure 1D). Together, these results point to sensitivity in positioning of HARs within 3D genome architecture similar to evolutionarily recent mammal and primate genes, while UCEs maintain consistency in their functional associations in the same manner as evolutionarily older metazoan and chordate genes.

### The functional associations of regions of varying sequence conservation are more sensitive to disease state in loop anchors than in domains

Due to potential changes in chromatin folding and gene regulation in disease states^17,20,124^, we next asked whether healthy and diseased cancer states can impact the positioning and the functional associations of regions of varying sequence conservation within 3D genomic features. Correlations between pooled domains and metazoan, mammal, and primate genes as well as UCEs change from non-significant in one state to significant in the other (2.29×10^-7^ ≤ p ≤ 0.870; Figure 4A; Table S2). Despite this variation in 3D positional relationships, the functional associations of those regions seem mainly unaffected. Broadly, era genes, UCEs, and HARs in pooled healthy domains (Figures S7A-S7F) and pooled cancer domains (Figures S7G-S7L) are enriched for very similar GO terms, as seen in all pooled domains (Figures 3, S6K, and S6L). For pooled loop anchors, we observed changes in the correlations for chordate and mammal genes along with HARs between healthy and cancer states (Figure 4A). As for functional associations, although GO terms associated with era genes, UCEs, and HARs in pooled healthy loop anchors (Figures 4B and S8A-S8E) closely resemble those found with all pooled loop anchors (Figures S6A-S6F), GO term enrichments in pooled cancer loop anchors can show notable shifts depending on the class of region (Figures S8F-S8I). For instance, older metazoan and chordate genes and UCEs in pooled loop anchors have largely similar GO term enrichments in cancer (Figures S8G-S8I) and healthy (Figures S8A, S8B, and S8E) states, potentially suggesting the exploitation of signaling pathways by cancer cells^125^. In contrast, younger mammal and primate genes in pooled cancer loop anchors show no GO term enrichments. As for HARs in pooled loop anchors, we noted changes in GO term enrichments between healthy (Figure 4B) and cancer states (Figure 4C). While HARs in pooled loop anchors in both states are still associated with neuronal-related processes, the individual GO terms are often state-specific. These diverse associations of HARs are in line with their implications in human neurodevelopment and disease^126,127^. In summary, the functional associations of evolutionarily recent mammal and primate genes in pooled loop anchors seem to be lost in disease, suggesting that cancer cells may better exploit the stable functional associations of older genes and UCEs.

**Figure 4.**
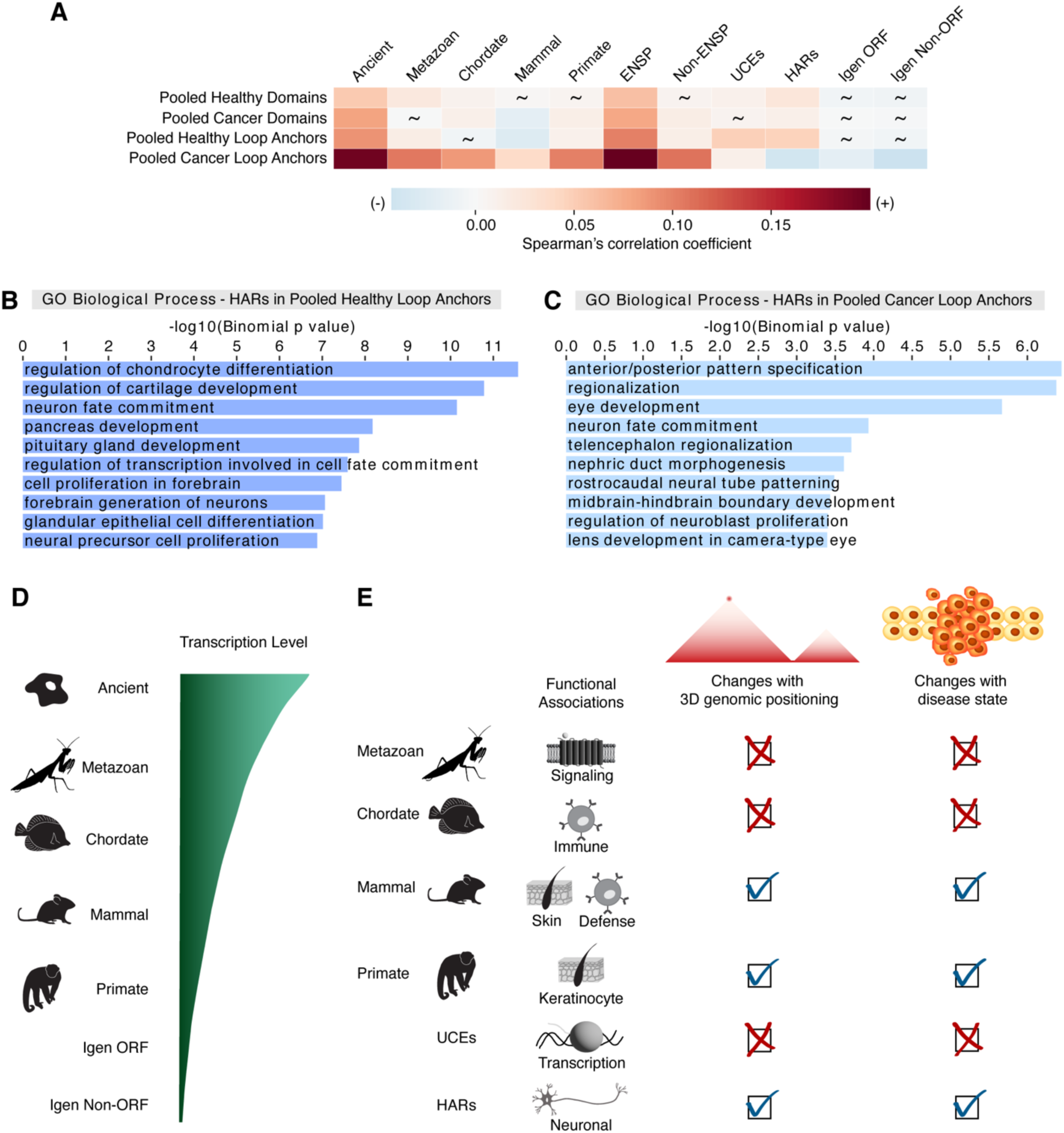
The relationship between genomic regions of different sequence conservation and 3D genome organization in healthy versus disease states. (A) Correlations of regions of varying sequence conservation with pooled domains and pooled loop anchors in healthy versus cancer states. Spearman correlation analysis was performed by partitioning the genome into 50-kb bins. Spearman correlation coefficients indicated by a heatmap. Non-significant p values are depicted by tildes. Control datasets include annotated genes, unannotated genes, Igen ORFs, and Igen Non-ORFs. (B and C) HARs that overlap pooled loop anchors in (B) healthy (dark blue) and (C) cancer (light blue) states against whole genome background share associations with neuronal GO categories. However, such HARs in healthy state are enriched for GO terms associated with cartilage, pancreas, and gland development, while HARs in cancer are associated with pattern specification, eye, and kidney development. (D) Evolutionary progression of transcription levels: RNA expression increases with time from youngest genes (primate) to oldest genes (ancient) and is higher in genes than in control non-genic sequences (Igen ORF, Igen Non-ORF). (E) Schematic of the functional associations of genomic regions of varying sequence conservation and their changes with 3D genomic positioning and disease state. Change is indicated by a check mark and consistency is depicted by an X.

## Discussion

Our study reveals that the relationships between genes of different evolutionary ages and their positioning within domains and loop anchors are variable, while the correlations of those era genes with boundaries display remarkable consistency across cell types. The transcription levels of genes also increase with evolutionary age in all tissue categories (Figure 4D). As for the functional associations of era genes, they are predominantly related to distinct cell type-specific processes. The functional associations of more recently evolved genes as well as HARs are sensitive to their 3D genomic positioning. This sensitivity is especially prominent in loop anchors compared to domains in disease state (Figure 4E). Loop formation has been implicated in demarcating domains and facilitating contacts between enhancers and promoters^6,9,12^. Although domain and loop structures may be changing, at least half may be preserved across cell types^5,6,128^. Ancient genes exhibit more consistent positive correlations with both domains and loop anchors between cell types, while less conserved, younger genes, with their marked increase in correlation variability, may be linked to cell type-specific chromatin organization. Furthermore, evolutionarily younger genes are associated with cell type-specific functions, suggesting their potential role in tissue-specific programs. Interestingly, previous studies have shown that many loops involving enhancers and promoters may be tissue-specific during differentiation, and therefore related to cell type-specific gene regulation^129,130^. Hence, the relationships of these recent era genes with 3D genomic features may be implicated in the evolution of tissue-specific genome activity. Moreover, modifications to the dynamic 3D regulatory landscape may even shape species-specific programs^131–136^.

Notably, analyzing transcriptional levels from GTEx shows an evolutionary progression of gene expression: young genes are expressed at lower levels than ancient genes, yet at much higher levels than background genomic transcription. This evolutionary progression of gene expression is observed with two different gene age estimation methods, synteny and sequence similarity. The output of gene regulation by way of various gene regulatory elements, including 3D genome organization elements, is gene expression. The placement of 3D genome organization elements may be used to control the access of transcriptional complexes to DNA, which may then produce RNA. Interestingly, for genes of all ages in the six tissue categories we considered, RNA levels are highest in testis, a tissue where DNA is loosely packed with protamines instead of histones^137^, and are unexpectedly lowest in the brain, where gene expression is tightly regulated but there are many cell types. Indeed, some brain areas such as the cerebellum do have high RNA expression. The similar evolutionary progression of RNA levels in all tissues suggests RNA levels are regulated by the 3D genome organization elements implicated in opening chromatin, and thus in part not by cell type-specific transcription factors – while a few transcription factors, like CTCF^22^, change genome organization, most transcription factors regulate expression differently, by cooperatively aiding the assembly of the RNA polymerase II transcriptional complex for particular genes. These two facets of gene regulation support transcription of a new gene and are stabilized evolutionarily if the gene finds a function and perdures evolutionarily. Additionally, longer genes of all ages are expressed at higher levels than shorter genes of all ages, suggesting gene size, usually higher in neuronal genes^82,138^, is associated to gene expression level independently of gene age.

The functional associations found with genes of each era are intriguing in light of the biological processes that arose at different times in evolution^80,117^. In line with this notion, the SMAD-dependent and G-protein coupled receptor signaling pathways associated with metazoan genes are often involved in altering gene expression related to tissue differentiation, which is critical for development of multicellular animals^139^. Chordate genes as well as mammal genes are linked to several immune processes, which may mediate defense responses against pathogens^140,141^. Both mammal and primate genes, recently shown to be associated with genetic individuality^142^, are associated with keratin processes, possibly related to hair formation, which may be integral to heat retention and homeothermy^143^. Interestingly, these links to immune response and skin development are further supported by the functional associations of fast-evolving genes and regulatory landscape in mammals facilitating an organism’s interactions with the environment^51,144^. Overall, the emergence of genes at different evolutionary times may be linked to novel biological processes gained at different levels of the phylogenetic hierarchy^80,117^.

Disruptions of the 3D genome architecture can play roles in various diseases, especially when interactions between regulatory regions are affected^13,14,17,20,129^. In particular, the identity and evolutionary age of the genes impacted by such alterations may be integral to understanding a disease. For instance, previous reports indicate a significant over-representation of disease-associated genes, including cancer-associated genes, that predate or originated with the emergence of Metazoa^117,125^. Interestingly, we observed little change in the functions associated with metazoan genes in loop anchors between healthy and cancer states, possibly related to the exploitation of signaling pathways by cancer cells in multicellular organisms^125^. As for evolutionarily recent mammal and primate genes in loop anchors, both lose all functional GO term enrichments in cancer, consistent with the under-representation of such genes among cancer and other disease-associated genes^117,125^. Nevertheless, fast-evolving HARs in loop anchors maintain functional associations with neuronal processes in both healthy and cancer states, reminiscent of the other disease implications of HARs, including in neurological disease^126,127^.

In conclusion, we describe the functional significance of the 3D genomic positioning of genes of different evolutionary ages and lineage restriction levels. The evolution of genes and their regulatory regions within the 3D genome may present different lenses through which the determinants of yet unresolved genetic diseases may be elucidated.

## Materials and Methods

### Human sequence datasets

As in other studies^82,145^, we retrieved 19,334 human annotated proteins from the Ensembl 89 database (www.ensembl.org), keeping the oldest and longest isoform per protein-coding gene. Additionally, we used 2,102 human unannotated protein sequences that are proteogenomically confirmed^76–79^. Furthermore, we used two kinds of control sequences. The first are Igen ORFs, localized in the non-genic non-repetitive part of the human genome, starting with a methionine (Met) codon and having at least 40 non-STOP codons, followed by a STOP codon. These sequences have no known evidence of transcription or translation but could in principle encode proteins, based on their DNA sequence. The second type of controls are intergenic DNA sequences, randomly chosen from the non-repetitive part of the human genome, which do not overlap with any Igen ORFs or known protein-coding genes, do not start with Met, may have STOP codons and are not expected to encode open reading frames (Igen Non-ORFs). Igen Non-ORFs were chosen to have the same DNA sequence length distribution as Igen ORFs.

### Gene age estimation by sequence similarity

We estimated the evolutionary age of protein-coding genes by sequence similarity^80,81^, which classifies genes based on their phylogenetic lineage restriction as estimated by protein sequence similarity^82^. Sequence similarity was evaluated with BLASTP^146^ and in some cases followed by HMMER^147^ to find the most taxonomically distant species in which a sufficiently similar protein sequence exists, such that the e-value of that most distant BLASTP hit is less that 10^-3^ ^82^.

#### 1. Building the dataset for gene age estimation

i. Protein sequences were downloaded from Ensembl version 89. 80,421 proteins (for 19,571 genes) were coming from the “canonical” chromosomes 1-22, X, and Y.
ii. 2,624 unannotated proteins were added^76–79,148^.
iii. Protein length filtering: Proteins < 40 amino acids (AA) and ≥ 4,000 AA were removed. First, lineage restriction levels cannot be reliably assigned for short proteins (< 40 AA) that may have evolved multiple times. Extremely long proteins (≥ 4,000 AA) did not run properly on certain tools in the pipeline.

Applying steps (i-iii), the resulting set of annotated and unannotated proteins had 65,898 proteins.

#### 2. Gene age estimation was performed by determining the taxonomic restriction levels of 65,898 proteins, assigning each to a phylostratum (PS) of 1-31

i. Lineage restriction determination. For most species, the NCBI Taxonomy database indicates the successive divergence nodes in the lineage of a species (for example, https://www.ncbi.nlm.nih.gov/Taxonomy/Browser/wwwtax.cgi?mode=Info&id=9606&lvl=3&lin=f&keep=1&srchmode=1&unlock). The intervals between successive nodes are termed phylostrata. Within the human lineage, there are 31 taxonomic nodes, yielding to 31 phylostrata (NCBI Taxonomy, https://www.ncbi.nlm.nih.gov/Taxonomy/Browser/wwwtax.cgi?id=9606). Node evolutionary timing was estimated using TimeTree^149^. Minimal gene age was assessed as the time when the most recent common ancestor (of the query species and the most distant species in which a sufficiently similar protein sequence was found) lived.
ii. BLASTP (v.2.6.0+)^146^ was used to evaluate protein sequence similarity. Each protein sequence was used to query the NCBI NR database (February 2017).
iii. Significance levels: an e-value limit of .001 was used to avoid false positive hits.
iv. Within input protein sequences, low-complexity sequence, which could yield chance hits^150^, was masked with SEG^151^.
v. Search length: up to 200,000 hits were allowed for each protein. Otherwise, the search could end before identifying the most distant hit and result in incorrect lineage restriction assignments.
vi. After assigning a taxonomic restriction level and thus a minimal evolutionary age to each protein, the longest of the oldest isoforms was chosen for each protein.

- For Ensembl-annotated proteins, proteins with the same ENSG ID were considered isoforms.
- For other proteins, isoforms were assigned based on chromosomal overlap.
- This approach yielded 19,334 annotated and 2,102 unannotated proteins, for a total of 21,436.
vii. Taxonomic restriction groups. The 31 phylostrata were grouped into 5 large-scale evolutionary eras: Ancient (PS 1-3), Metazoan (PS 4-7), Chordate (PS 8-17), Mammal (PS 18-22), Primate (PS 23-31).
viii. Technical constraints considered for fusing individual nodes into large-scale evolutionary eras:

- Since for some phylostrata there are only one or few outgroup species with genomes that are well-sequenced and annotated, gene loss could lead to a misassignment by 1-2 phylostrata.
- For some phylostrata, the interval between successive divergence times is estimated by TimeTree^149^ to have a duration of 0 My. Thus, such phylostrata cannot be separated and should be fused.
- Some phylostrata have small numbers of genes.

Given these constraints, more robust conclusions can be drawn for larger batches of genes, with broader ranges of taxonomic restriction assignments, and thus broader ages.

### Comparison between sequence similarity and synteny for gene age estimation

#### 1. Comparison procedure

We compared the lineage restriction levels obtained by chromosomal synteny^83,152^ or by protein sequence similarity. For every gene, we used the lineage restriction level obtained with synteny-based methods in the GenTree database^83^ and compared it with the lineage restriction level obtained with sequence similarity in the 18,098 human genes common to the GenTree database and to our human gene list. Since GenTree only includes Ensembl-annotated genes, the common gene list does not include the unannotated genes, most of which appeared evolutionarily recently (Figure S1B).

#### 2. Gene dataset organization and quality control filters

To ensure reproducibility and consistency of gene counts and RNA transcript mapping, it is important to list for every protein-coding gene the complete protein sequence, gene name and description, chromosomal coordinates and ensure clean mapping in GRCh37 and GRCh38 (Table S1). With these criteria in mind, 18,098 (93.6% of our 19,334 annotated human genes) were retained as common between the GenTree database (which uses gene names) and our list.

#### 3. Taxonomic restriction groups used for comparison

To compare the results of synteny and sequence similarity, we grouped the 31 human taxonomic nodes into 4 evolutionary eras: Early (PS 1-13), Vertebrate (PS 14-17), Mammal (PS 18-22), Primate (PS 23-31). We used this grouping since GenTree^83^ uses a subset of 15 of the 31 human taxonomic nodes, such that their oldest group (“branch”) encompasses all nodes from PS 1 (Cellular organisms) to PS 13 (Euteleostomi). Accordingly, this constrains the boundaries of the first two eras (Early, Vertebrate).

#### 4. Alternative methods considered

Of note, to evaluate gene lineage restriction and thus gene age, we did not rely on gene orthology. The reason is that different rates of gene duplication and of gene loss between lineages, as well as large-scale chromosomal recombination, both lead to a loss of 1:1 correspondence between similar genes in different species and to a loss of synteny. Thus, the number of orthologs confirmed by synteny decreases rapidly with evolutionary distance^90^.

### UCE and HAR datasets

The UCE dataset includes 896 elements from comparisons between human-mouse-rat, human-dog-mouse, or human-chicken genomes as previously reported^39,53,91^. The genomic coordinates for HARs were obtained from Girskis *et al.*^43^ and converted to the hg19 genome assembly using the UCSC LiftOver tool (https://genome.ucsc.edu/cgi-bin/hgLiftOver). Any overlapping intervals were collapsed to avoid including the same region more than once.

### Hi-C datasets

The coordinates for Hi-C annotated genomic features, including domains, boundaries, and loop anchors, were retrieved from published datasets as indicated in Table S1^5,6,19,93–103^. The genomic coordinates were processed as previously described^39^. Namely, when applicable, the Hi-C genomic annotations were lifted over to the hg19 genome assembly using LiftOver. To ensure that any given region was counted only once, overlapping intervals were collapsed to generate final pooled and individual datasets for domains, boundaries, and loop anchors that may differ from previously published original ones. Information about these final datasets, including number of regions, median region size (bp), coverage (bp), and proportion of genome covered (%), is provided in Table S1. Eres *et al.*^95^ provided four iPSC datasets derived from four individuals, while Dixon *et al.*^5^ and Rao *et al.*^6^ each provided one dataset of IMR90 using dilution and in situ Hi-C protocols, respectively. Rubin *et al.*^100^ provided three Hi-C datasets corresponding to different stages of keratinocyte differentiation (days 0, 3, and 6). Dixon *et al.* 2012^5^ and Dixon *et al.* 2015^94^ each provided a Hi-C dataset for H1 human embryonic stem cells. Due to the methodological variations among these studies such as Hi-C protocol, quantity of input material, and sequencing depth, we examined each dataset individually, grouped by a Hi-C feature (e.g. Pooled Domains), and grouped additionally based on healthy or disease state (e.g. Pooled Cancer Domains).

### Correlation analyses between 3D genomic features and regions of varying sequence conservation

Correlation analyses were performed using a previously established method^39,53^. Briefly, the genome was first split into bins of fixed size. In each bin, the proportion of sequence covered by a 3D genomic feature, or regions of varying sequence conservation was calculated. Then, for each bin the global correlations among those feature densities of interest were determined; for example, the densities of primate genes and Pooled Domains were correlated across the same binning of the genome. The Spearman correlation coefficients and the corresponding p values were reported for a pairwise comparison between two features (e.g. the pairwise correlation between primate genes and Pooled Domains) using three different bin sizes (20 kb, 50 kb, 100 kb; Table S2). The Spearman correlation coefficients were also indicated by a heatmap.

### GTEx analysis

To determine the level of RNA expression of all human protein-coding genes encoding proteins of at least 40 AA and of control intergenic sequences, an RNA-Seq calling pipeline was used to calculate the number of reads per gene per sample broken down by tissue. GTF files with all sequences (annotated and unannotated genes, Igen controls) were run through the GTEx RNA-Seq calling pipeline of the Broad Institute (https://github.com/broadinstitute/gtex-pipeline). Deseq2 normalization of means was used (https://github.com/broadinstitute/pyqtl) to account for sample-level sequencing depth between individuals and RNA composition. From this, we obtained a mean read count value for each sequence for each of the 54 tissues in GTEx. These tissues were aggregated into the following tissue categories: three germ layers (ectoderm, mesoderm, endoderm), two germline tissues (ovary, testis) and one tissue of special interest to the authors (brain). To avoid overweighting brain regions in ectoderm, only cortex and cerebellum were included (and these tissues were also included in brain, along with 11 other brain regions). Similar analyses were conducted with transcripts per million (TPMs) per gene, as available from the online resource of GTEx for annotated genes alone (v8, median TPM counts per gene). TPM data was downloaded directly from the GTEx v8 Browser’s download page (https://storage.googleapis.com/adult-gtex/bulk-gex/v8/rna-seq/GTEx_Analysis_2017-06-05_v8_RNASeQCv1.1.9_gene_median_tpm.gct.gz), with median TPM counts per tissue, calculated from 54 tissues for every annotated gene. The purpose of the TPM analysis was checking if the changes with evolutionary age observed with Mean Counts remain the same with TPMs, which better accounts for transcript length but does not work as well for small genes.

Transcripts of human genes encoding proteins of at least 40 AA (19,334 annotated protein-coding genes, 2,102 unannotated proteogenomically confirmed genes) and control sequences were counted in 54 human adult tissues (GTEx database) using GRCh38 genomic coordinates for all sequences. Of 20,000 randomly picked Igen ORFs (ten sets of 2,000 each), for our analysis we used two sets of 2,000 each (4,000 sequences total), from which 3,916 Igen ORFs passed GTEx filtering consisting of not duplicating or overlapping other Igen ORFs, and cleanly mapping to GRCh37. Of 20,000 randomly picked Igen Non-ORFs, for our analysis we used two sets of 2,000 each (4,000 sequences total), from which we used 3,889 Igen Non-ORFs that were filtered to not include duplicates; not overlap other control Igen ORFs, Igen Non-ORFs, or genes; and map cleanly to GRCh37. The same approach was applied to 20,000 GRCh37 Igen ORFs and 20,000 Igen Non-ORFs, resulting in 3,924 Igen ORFs and 3,892 Igen Non-ORFs, which cleanly mapped to GRCh38 and passed the filtering described previously. A small number of genes (less than 1.3% of all annotated genes) do not map fully between GRCh37 and GRCh38 (247 genes with the same age distribution as all genes; 179 ancient, 32 metazoan, 25 chordate, 8 mammal, 3 primate). The data were analyzed as a function of gene evolutionary age. Statistical comparisons between gene age categories (genes of different eras and the controls) and between tissue groups were done with the Mann-Whitney U test, with p values corrected for multiple comparisons with the Benjamini-Hochberg correction. P values of 0 indicate numerical values smaller than 2.225×10^-308^. Data visualization was done using the Python library, Seaborn 0.12.2 (https://github.com/mwaskom/seaborn/tree/v0.12). Boxes are from the 25^th^ to the 75^th^ percentile; medians are indicated by horizontal lines; whiskers indicate 1.5 times the interquartile range, with outliers not shown (but included in analysis). For visualization purposes, we applied a log10 data transformation, and handled 0.00 values by adding the smallest non-zero value to every log10 argument.

### GREAT analysis

Genomic Regions Enrichment of Annotations Tool (GREAT, version 4.0.4)^118^ analysis was performed to assess gene ontology (GO) terms associated with genes in each era, UCEs, or HARs. This approach of using genomic coordinates of interest allowed inspection of HARs, UCEs, and yet unannotated novel genes, especially in the category of primate genes, that may have otherwise gone amiss with more conventional approaches that rely on gene names or identifiers as inputs^118^. Furthermore, subsets of regions of varying sequence conservation that overlap with 3D genomic feature of interest were determined using pybedtools wrapped bedtools intersect^153,154^. The argument u=True ensured that the same region was written only once if any overlaps were found with 3D genomic feature. Such regions that intersected a 3D genomic feature were assessed for their association with GO terms against the relevant background. For genes in each era, the total list of all era genes was used as a background. In the case of UCEs and HARs, the whole genome provided by GREAT was used as a background.

## Data Availability

The publicly available datasets were obtained using accession numbers GSE71831^93^, GSE71072^96^, GSE63525^6^, GSE77565^103^, the 3D Genome Browser (3dgenome.org), dbGaP (GTEx, phs000424.v8.p2), and publications as provided in the Table S1.

## Acknowledgments

We are grateful to C.-ting Wu and Marc Kirschner for discussions. We thank the Computational Biology Core within the Institute for Systems Genomics at the University of Connecticut for access to the high-performance computing cluster resources. We apologize to the authors whose work we could not include due to space constraints. N.G. was supported by a Summer Undergraduate Research Fund (SURF) award. Work in J.E.’s laboratory was supported by the University of Connecticut and an award to J.E. from NIH/NIGMS (R35GM146922). V.L. and N.S. were supported by NIH grants R01NS095654, P50MH106934, U01MH116488, and R01MH110926

(N.S.), and the Simons Foundation (N.S.). D.M., W.P., and A.O.D.L. were supported by the Manton Center for Orphan Disease Research Scholar award (A.O.D.L.).

## Author Contributions

K.F., V.L., N.S., A.O.D.L., and J.E. designed research. K.F., V.L., N.G., A.K., T.H., D.M., W.P., K.-M.N., N.S., A.O.D.L., and J.E. performed research and analyzed data. K.F., V.L., and J.E. wrote the paper. All authors approved the final version of this manuscript.

## Supplemental information

**Figure S1.**
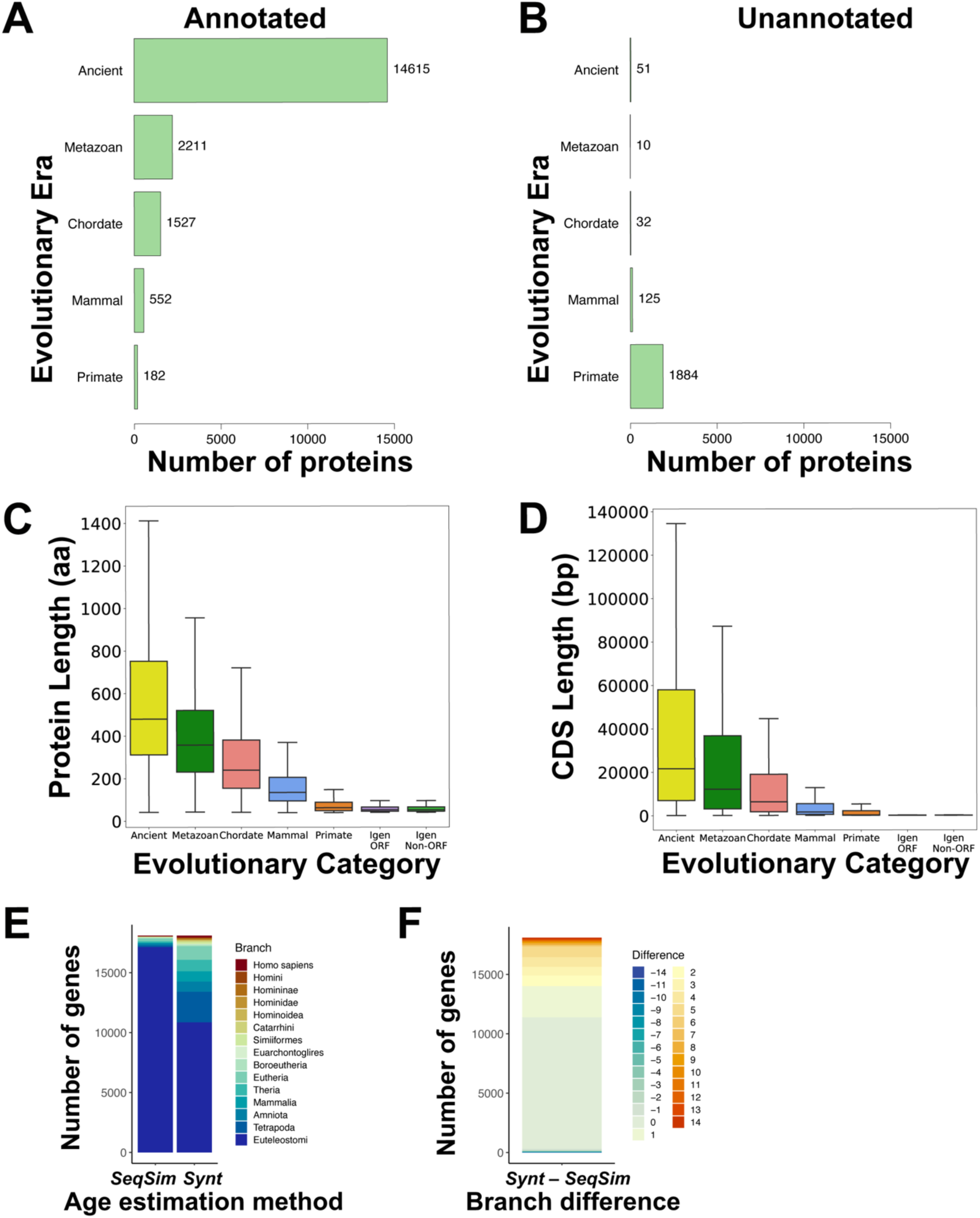
Gene numbers and lengths for each evolutionary era. (A) Number of annotated human protein-coding genes in every evolutionary era (ancient, metazoan, chordate, mammal, primate). Most annotated genes are evolutionarily ancient. (B) Number of unannotated human protein-coding genes in every evolutionary era (ancient, metazoan, chordate, mammal, primate). Most unannotated genes are evolutionarily young. (C) Protein length is indicated in number of amino acids (AA). The evolutionarily youngest genes are much shorter than ancient genes and only slightly longer than control Igen ORF and Igen Non-ORF sequences. (D) The length of coding sequences (CDS) is indicated as the number of base pairs (bp). The evolutionarily youngest genes are shorter than ancient genes and similar to control Igen ORF and Igen Non-ORF sequences. (E-F) Comparison of gene age estimated by sequence similarity (SeqSim) and synteny (Synt) in 18,098 human genes. (E) The number of human protein-coding genes is indicated in 15 evolutionary branches used in the GenTree synteny-based database^1^. More genes appear evolutionary young by synteny than by sequence similarity. (F) Difference between the branch number of every gene as evaluated by synteny or sequence similarity. Most genes show no difference. A substantial fraction appears younger by synteny. Very few appear younger by sequence similarity.

**Figure S2.**
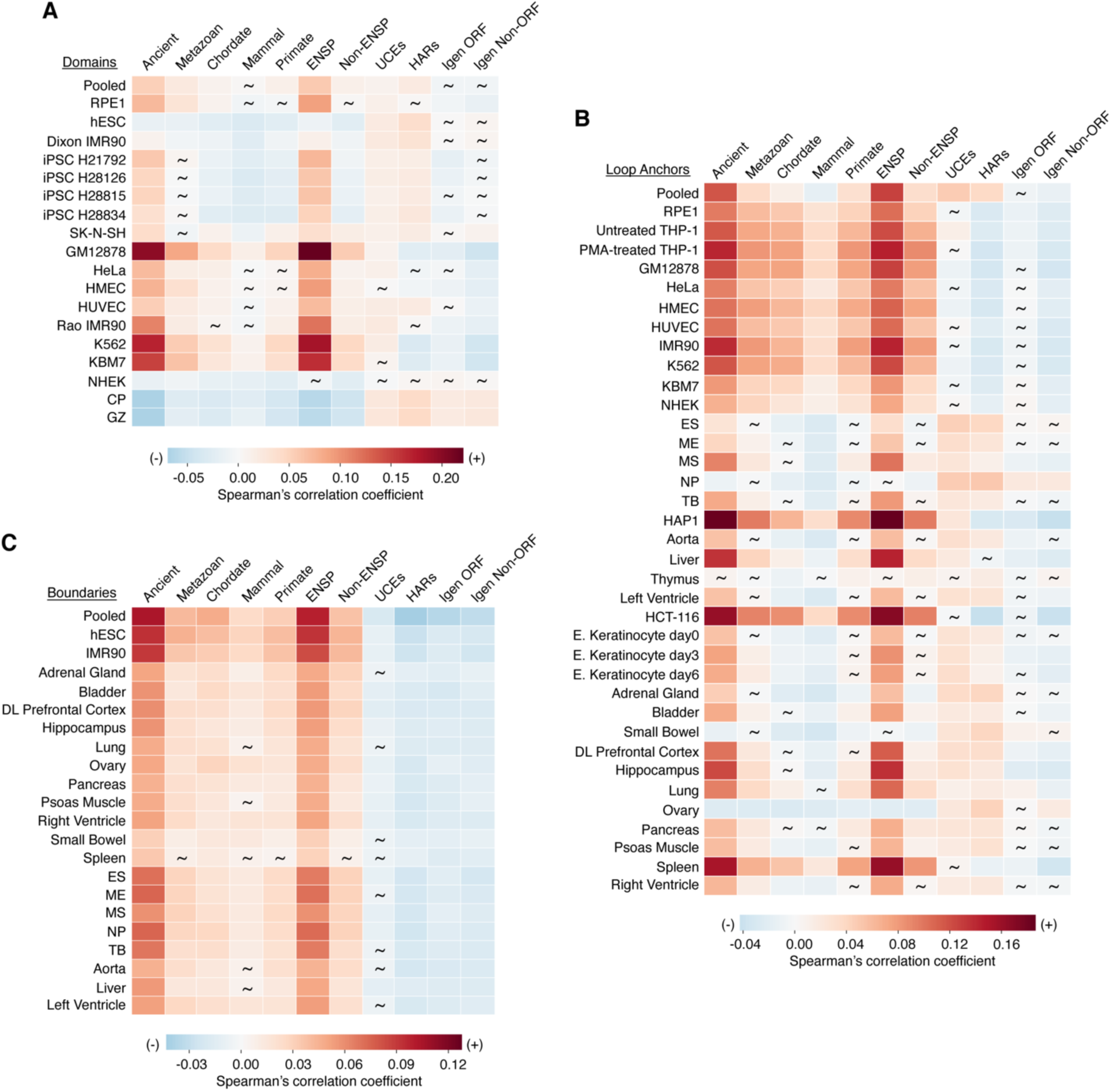
The relationships between 3D genomic features and various evolutionary genomic regions show stability or variation across cell types. (A) Ancient genes, annotated genes, and UCEs are primarily significantly positively correlated with domains (0 ≤ p ≤ 0.746). Conversely, less conserved genes and fast-evolving HARs show variable relationships with domains across cell types (2.34×10^-92^ ≤ p ≤ 0.988). (B) The relationships between loop anchors and regions of various sequence conservation show different variation levels across cell types. Ancient and annotated genes have more consistent relationships with loop anchors by cell type (0 ≤ p ≤ 0.944), whereas less conserved genes, HARs, and UCEs have more variable relationships (8.65×10^-118^ ≤ p ≤ 0.965). (C) The relationship of boundaries with era genes, UCEs, and HARs is highly consistent across cell types. Era genes are predominantly significantly positively correlated with boundaries (2.19×10^-110^ ≤ p ≤ 0.306), while UCEs and HARs are non-significantly or significantly negatively correlated (8.91×10^-12^ ≤ p ≤ 0.136). (A-C) Spearman correlation analysis was performed by partitioning the genome into 50-kb bins. Spearman correlation coefficients are indicated by a heatmap. Non-significant p values are depicted by tildes. Control datasets include annotated genes, unannotated genes, Igen ORFs, and Igen Non-ORFs. CP, cortical and subcortical plate; GZ, germinal zone; E., epidermal; DL, dorsolateral.

**Figure S3.**
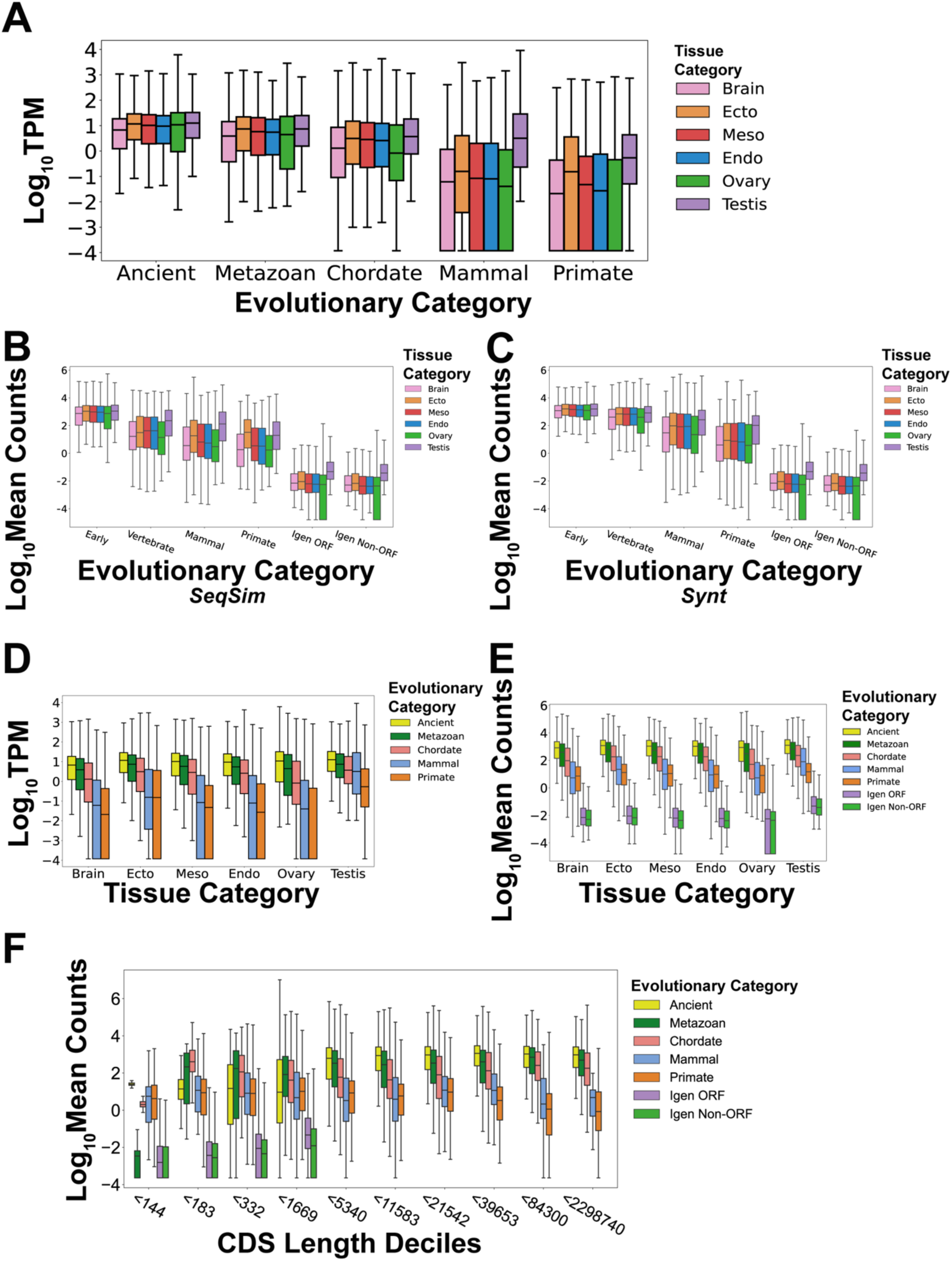
Transcriptional level variation with gene age, tissue of origin and gene length. As in Figure 2, the expression of human protein-coding genes is measured for RNA transcripts across 54 human tissues from the GTEx database^2^. (A) RNA expression levels, measured as log10 of TPMs, increase from youngest to oldest genes (Table S3C). Within mammal and primate genes, expression in testis is higher than in other tissues. (B-C) RNA expression of human genes of different evolutionary ages as a function of the gene age estimation method – sequence similarity, SeqSim (B), synteny, Synt (C). Genes of different lineage restriction levels, or gene ages, are grouped into 4 eras (early, vertebrate, mammal, primate) that accommodate the 15 evolutionary branches used by GenTree^1^. Data are normalized counts of RNAs across 54 human tissues from the GTEx database^2^. The evolutionary change from low levels in young genes to high levels in early genes is very similar between the two methods. (D) RNA expression levels, measured as log10 of TPMs, in 6 tissue categories increase from youngest genes to oldest genes. (E) RNA expression levels in 6 tissue categories: brain, 3 germ layers (ectoderm, mesoderm, endoderm), and 2 germline tissues (ovary, testis). Within every tissue category, expression increases from youngest genes to oldest genes and is higher in genes than in control non-genic sequences. (F) RNA expression levels, measured as log10 of mean counts, increase with the gene length, shown as CDS length. Within 7 CDS length deciles, ancient genes are expressed at highest levels while primate genes are expressed at lowest levels. Within 3 of the shorter CDS length deciles, the highest expression is that of metazoan genes or chordate genes. Only the first 4 deciles have control non-genic sequences, and their expression is 100-1000 times lower than that of genes.

**Figure S4.**
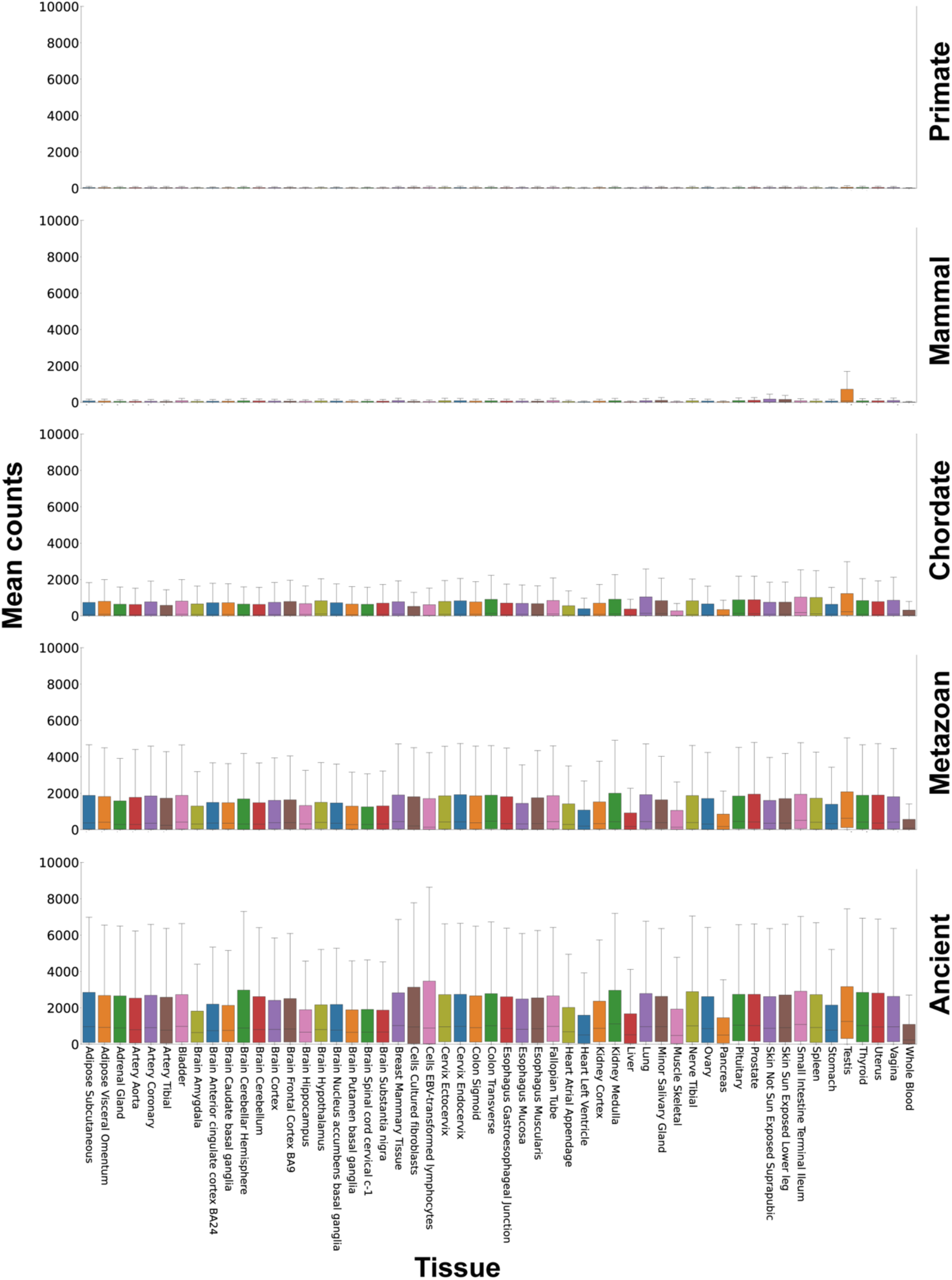
Gene expression in 54 human tissues and evolutionary eras. Gene expression levels are measured as mean counts of RNA transcripts across 54 human tissues in 5 evolutionary eras, as in Figures 2 and S3. Gene expression increases evolutionarily from primate to ancient genes. With each evolutionary era, testis expression is typically highest, followed by other tissues which differ between eras.

**Figure S5.**
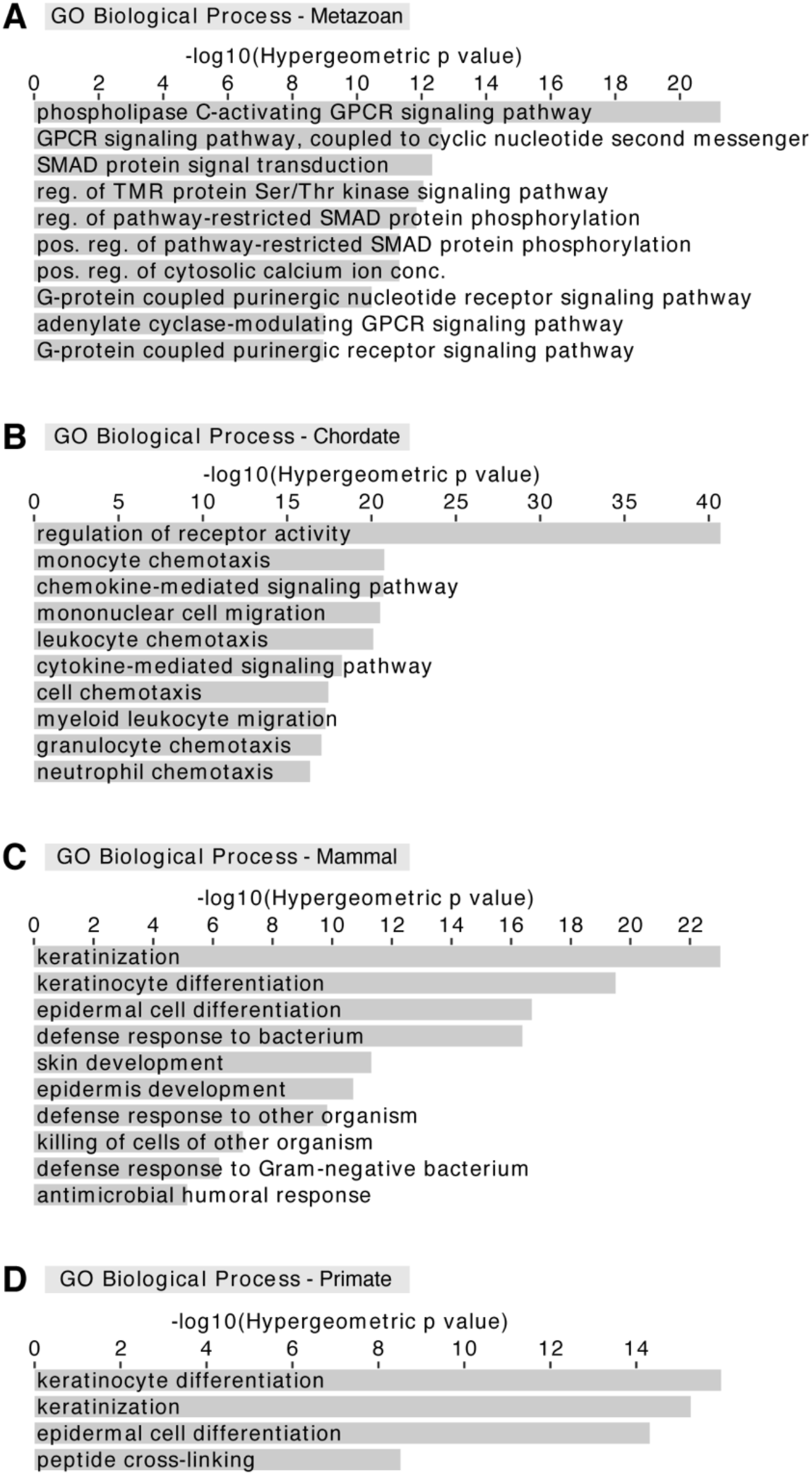
Functional associations of era genes. GREAT analysis for (A) metazoan genes, (B) chordate genes, (C) mammal genes, and (D) primate genes. Those GO terms are associated with very similar processes as for era genes in pooled domains (Figure 3). (A-D) Analysis performed against all era genes as a background. Abbreviated GO terms: GPCR, G-protein coupled receptor; reg., regulation; TMR, transmembrane receptor; Ser, serine; Thr, threonine; pos., positive; conc., concentration.

**Figure S6.**
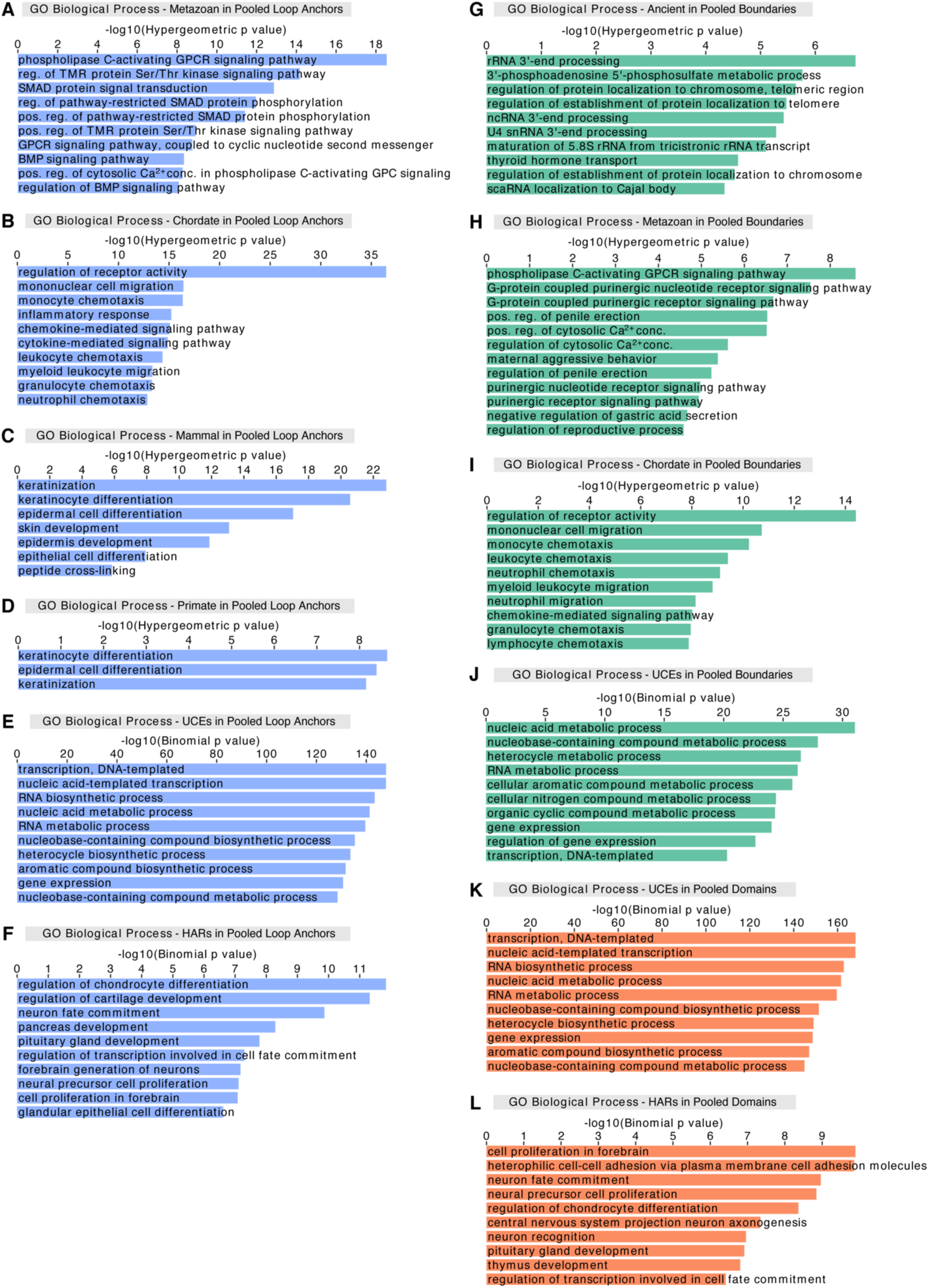
Functional associations of era genes, UCEs, and HARs within 3D genomic features. Regions of varying sequence conservation overlapping pooled loop anchors are linked to GO terms associated with (A) signaling pathways for metazoan genes, (B) immune response-related processes for chordate genes, (C) skin development for mammal genes (note absence of defense response compared to Figure 3C), (D) keratinocyte differentiation for primate genes (note absence of peptide cross-linking compared to Figure 3D), (E) transcription and RNA metabolic processes for UCEs, and (F) neuronal processes, cartilage formation, pancreas development, and glandular epithelial cell differentiation for HARs. In pooled boundaries, enrichments for GO terms are related to (G) protein localization to telomeres and ncRNA processes for ancient genes, (H) signaling pathways and regulation of reproductive processes for metazoan genes, (I) immune response-related processes for chordate genes, and (J) transcription and RNA metabolic processes for UCEs. (K) UCEs in pooled domains are associated with transcription and nucleic acid metabolism, and (L) HARs in pooled domains are associated with neuronal processes, cartilage formation, and thymus development. Backgrounds: all era genes for individual eras of genes; whole genome for UCEs and HARs. Abbreviated GO terms: GPCR, G-protein coupled receptor; reg., regulation; TMR, transmembrane receptor; Ser, serine; Thr, threonine; pos., positive; conc., concentration; tricistronic rRNA transcript, tricistronic rRNA transcript (SSU-rRNA, 5.8S rRNA, LSU-rRNA). Blue, pooled loop anchors; green, pooled boundaries; orange, pooled domains.

**Figure S7.**
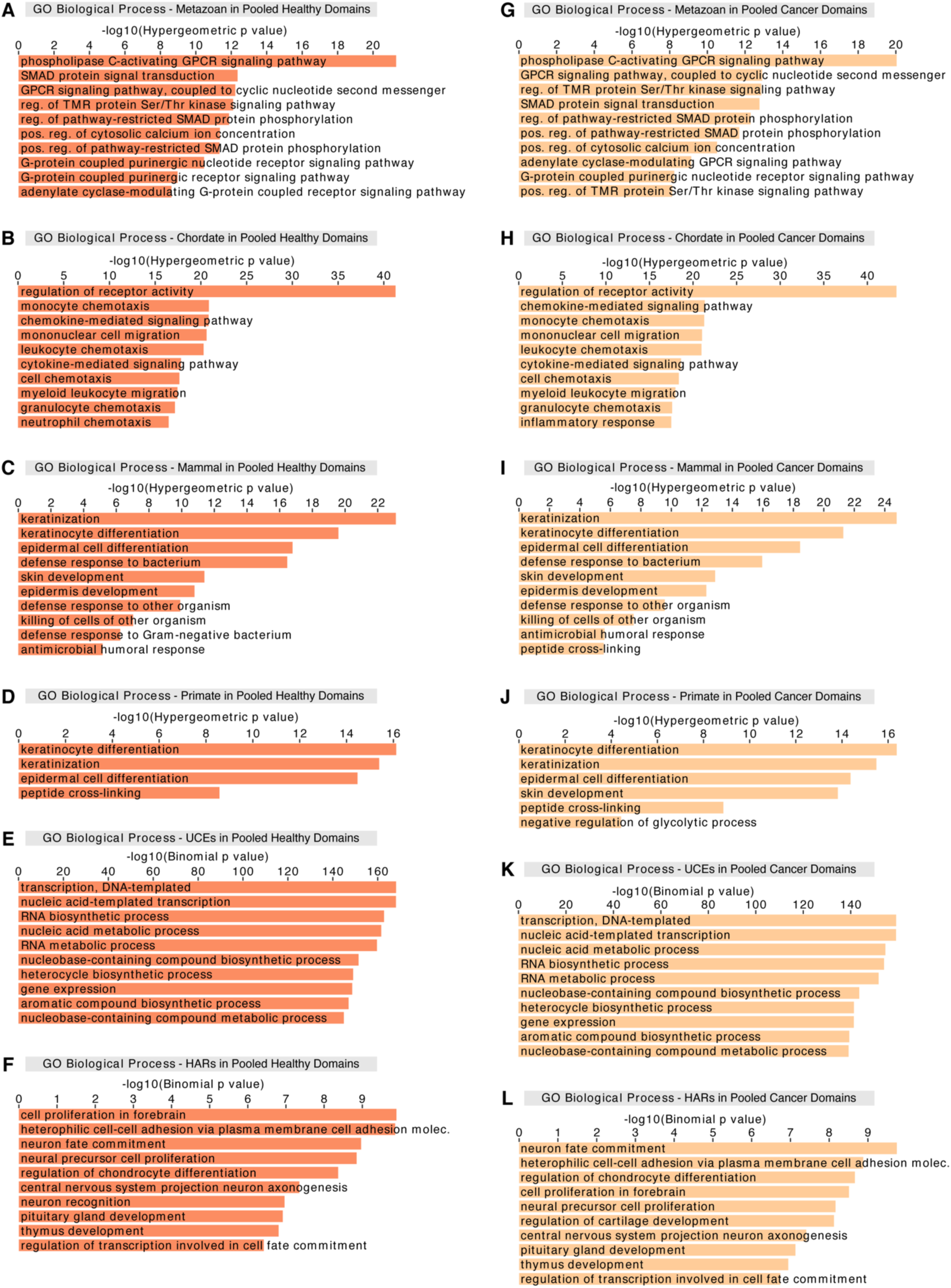
Functional associations of era genes in pooled domains in cancer state are very similar to those in all of the pooled domains and pooled healthy domains. (A-F) Era genes, UCEs, and HARs intersected with pooled domains in healthy state are associated with nearly identical processes as with all pooled domains (Figures 3, S6K, and S6L). (G-L) When intersected with pooled cancer domains, GO terms are very similar as those in all pooled domains (Figures 3, S6K, and S6L) and pooled healthy domains, with the exception of gained association with the glycolytic process in primate genes (J). GREAT analysis for regions of varying sequence conservation that overlap pooled healthy (A-F) and cancer (G-L) domains including (A and G) metazoan genes, (B and H) chordate genes, (C and I) mammal genes, (D and J) primate genes, (E and K) UCEs, and (F and L) HARs. Backgrounds: all era genes for individual eras of genes; whole genome for UCEs and HARs. Abbreviated GO terms: GPCR, G-protein coupled receptor; reg., regulation; TMR, transmembrane receptor; Ser, serine; Thr, threonine; pos., positive; molec., molecules. Dark orange, pooled healthy domains; light orange, pooled cancer domains.

**Figure S8.**
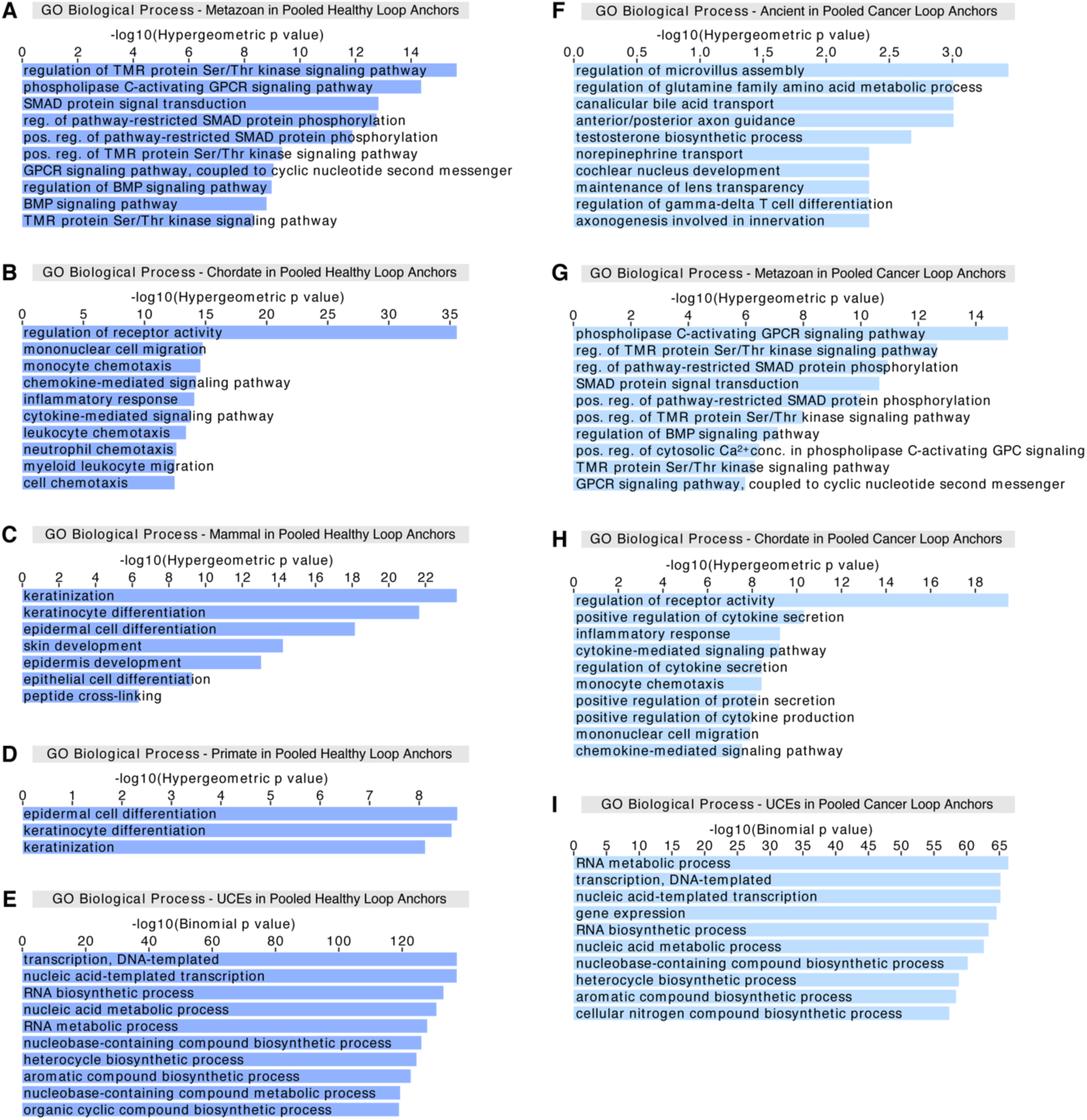
Evolutionarily older genes and UCEs exhibit GO term enrichments when in pooled cancer loop anchors. In pooled loop anchors in healthy state (dark blue), GO terms are associated with (A) signaling pathways for metazoan genes, (B) immune response-related processes for chordate genes, (C) skin development for mammal genes, (D) keratinocyte differentiation for primate genes, and (E) transcription and nucleic acid metabolism for UCEs. These GO terms are largely similar to those seen with all pooled loop anchors (Figures S6A-S6E). In pooled cancer loop anchors (light blue), GO terms are related to (F) a wide variety of processes for ancient genes, (G) signaling pathways for metazoan genes, (H) immune response-related processes for chordate genes, and (I) transcription and nucleic acid metabolism for UCEs. Note the absence of GO terms for mammal and primate genes. Backgrounds: all era genes for individual eras of genes; whole genome for UCEs. Abbreviated GO terms: TMR, transmembrane receptor; Ser, serine; Thr, threonine; GPCR, G-protein coupled receptor; reg., regulation; pos., positive; conc., concentration.

**Table S1.** Information on datasets.

(A) Information on era genes. (B) Control dataset for Igen ORF. (C) Control dataset for Igen Non-ORF. (D) Difference between genes ages estimated as lineage restriction levels by sequence similarity (SeqSim) versus synteny (Synt). (E) Hi-C datasets and related sources.

**Table S2.** Spearman correlation analyses of the relationships between regions of varying sequence conservation and 3D genomic features (domains, loop anchors, boundaries).

(A) Spearman correlation coefficients using 50-kb bins. (B) p values using 50-kb bins. Non-significant p values (red). (C) Spearman correlation coefficients using 20-kb bins. (D) p values using 20-kb bins. Non-significant p values (red). (E) Spearman correlation coefficients using 100-kb bins. (F) p values using 100-kb bins. Non-significant p values (red).

**Table S3**. Era genes and gene expression with taxonomic subdivision equivalence between NCBI nodes and GenTree branches.

(A) Mean counts of RNA transcripts in 6 adult human tissue categories in sequences from 7 evolutionary categories. There are 7 evolutionary categories of sequences: 5 evolutionary eras (Ancient, Metazoan, Chordate, Mammal, Primate), Igen ORFs, and Igen Non-ORFs. For every sequence, counts of RNA transcripts obtained using the GTEx database are shown, together with information about protein sequence, genomic localization, and gene evolutionary age. (B) Significance of statistical comparisons of mean counts from different tissue categories and evolutionary categories. Statistical significance was calculated with the Mann-Whitney U test and corrected for multiple comparisons using the Benjamini-Hochberg correction. (C) Counts of RNA TPM in 6 adult human tissue categories, in sequences from 7 evolutionary categories. As in (A), for every sequence, TPM counts from the GTEx database are shown, together with information about protein sequence, genomic localization, and gene evolutionary age. (D) Comparison of the expression of genes grouped by ages determined by sequence similarity versus synteny. Significance of statistical comparisons of mean counts from different tissue categories and evolutionary categories. Statistical significance was calculated with the Mann-Whitney U test and corrected for multiple comparisons using the Benjamini-Hochberg correction. (E) Significance of statistical comparisons of TPMs from different tissue categories and evolutionary categories. Statistical significance was calculated with the Mann-Whitney U test and corrected for multiple comparisons using the Benjamini-Hochberg correction. (F) Taxonomic subdivision equivalence between NCBI nodes (phylostrata, PS) and GenTree branches. The four evolutionary eras used for the comparison between Sequence Similarity and Synteny (Early, Vertebrate, Mammal, Primate) are indicated to the right. (G) For each protein-coding gene present in the GenTree database and in our Ensembl-derived database, the estimated evolutionary age corresponds to the taxonomic subdivision to which the gene belongs. The taxonomic subdivision is indicated as NCBI nodes (phyostrata, PS) and as GenTree branches. For the comparison between Sequence Similarity and Synteny, four evolutionary eras were used (Early, Vertebrate, Mammal, Primate) to accommodate the GenTree branches. For comparing counts between tissues, five evolutionary eras were used (Ancient, Metazoan, Chordate, Mammal, Primate). For each gene, indicated are the protein length, the amino acid sequence, and the differences between the age determined by synteny versus by sequence similarity.

